# Dissociation of the intramolecularly cleaved N- and C-terminal fragments of the adhesion G protein-coupled receptor GPR133 (ADGRD1) increases canonical signaling

**DOI:** 10.1101/2020.12.08.415398

**Authors:** Joshua D. Frenster, Gabriele Stephan, Niklas Ravn-Boess, Devin Bready, Jordan Wilcox, Bjoern Kieslich, Caroline Wilde, Norbert Sträter, Giselle R. Wiggin, Ines Liebscher, Torsten Schöneberg, Dimitris G. Placantonakis

## Abstract

GPR133 (ADGRD1), an adhesion G protein-coupled receptor (GPCR), is necessary for growth of glioblastoma (GBM), a brain malignancy. The extracellular N-terminus of GPR133 is thought to be autoproteolytically cleaved into an N-terminal and a C-terminal fragment (NTF and CTF). Nevertheless, the role of this cleavage in receptor activation remains unclear. Here, we show that the wild-type (WT) receptor is cleaved after protein synthesis and generates significantly more canonical signaling than an uncleavable point mutant (H543R) in patient-derived GBM cultures and HEK293T cells. However, the resulting NTF and CTF remain non-covalently bound until the receptor is trafficked to the plasma membrane, where we find NTF-CTF dissociation. Using a fusion of the hPAR1 receptor N-terminus and the CTF of GPR133, we demonstrate that thrombin-induced cleavage and shedding of the hPAR1 NTF increases receptor signaling. This study supports a model where dissociation of the NTF at the plasma membrane promotes GPR133 activation.

**Highlights:**

- GPR133 is intramolecularly cleaved in patient-derived GBM cultures
- Cleaved GPR133 signals at higher efficacy than the uncleavable GPR133 H543R mutant
- The N- and C-terminal fragments (NTF and CTF) of GPR133 dissociate at the plasma membrane
- Acute thrombin-induced cleavage of the human PAR1 NTF from the GPR133 CTF increases signaling

**eTOC Blurb:** Frenster et al. demonstrate intramolecular cleavage of the adhesion GPCR GPR133 in glioblastoma and HEK293T cells. The resulting N- and C-terminal fragments dissociate at the plasma membrane to increase canonical signaling. The findings suggest dissociation of GPR133’s N-terminus at the plasma membrane represents a major mechanism of receptor activation.

## INTRODUCTION

The adhesion family of G protein-coupled receptors (GPCRs) has attracted increasing interest in the recent years for essential functions in health and disease (Langenhan, 2019; Morgan et al., 2019). The large extracellular N-termini of adhesion GPCRs contain a GPCR-autoproteolysis inducing (GAIN) domain, which is thought to catalyze autoproteolytic cleavage at the GPCR proteolysis site (GPS) marked by the tripeptide sequence H-L/I-*-S/T (* denotes the cleavage site) (Arac et al., 2012; Lin et al., 2010). Following this intramolecular cleavage, adhesion GPCRs are generally believed to exist as non-covalently bound heterodimers of their extracellular N-terminal fragment (NTF) and transmembrane-spanning C-terminal fragment (CTF) (Gray et al., 1996; Krasnoperov et al., 1999). The recent demonstration of a tethered internal agonist, also known as the *Stachel* sequence, immediately C-terminal to the GPS, has given rise to the hypothesis that NTF-CTF dissociation facilitates the conformational changes needed for the *Stachel* sequence to initiate receptor activation (Liebscher et al., 2014).

However, the mechanism of receptor activation mediated by autoproteolytic cleavage and NTF-CTF dissociation is not generalizable to all members of the adhesion GPCR family. Indeed, several adhesion GPCRs have not yet been observed to undergo intramolecular cleavage and cleavage is not necessarily required for their activity (Bohnekamp and Schoneberg, 2011; Langenhan, 2019; Promel et al., 2012; Vallon and Essler, 2006; Wilde et al., 2016). Additionally, in some adhesion GPCRs cleavage occurs in selective cellular contexts but not others (Arac et al., 2012; Hsiao et al., 2009; Iguchi et al., 2008; Yang et al., 2017). Finally, cleavage- and *Stachel*-independent signaling have been reported for several adhesion GPCRs (Kishore et al., 2016; Patra et al., 2013; Promel et al., 2012; Salzman et al., 2017; Wilde et al., 2016). These observations emphasize the need to study mechanisms of activation for adhesion GPCRs on an individual basis and in physiologically relevant biological contexts.

We previously described that GPR133 (ADGRD1), a member of the adhesion family of GPCRs, is required for growth of glioblastoma (GBM), an aggressive primary brain malignancy (Bayin et al., 2016; Frenster et al., 2017; Frenster et al., 2020). In heterologous expression systems, N-terminally truncated CTF constructs of GPR133 generate significantly more G protein-mediated signaling than the full-length receptor (Liebscher et al., 2014). Nonetheless, there has not been any prior study of the extent of GPR133 cleavage or the NTF-CTF association. Here, we demonstrate that GPR133 is almost entirely cleaved in patient-derived GBM cells, and that cleaved GPR133 has a higher basal activity than an uncleavable GPR133 point mutant. While the cleaved CTF and NTF remain non-covalently bound to each other within the secretory pathway, we demonstrate that the NTF dissociates from the CTF once reaching the plasma membrane. Using a fusion protein of the N-terminus from human PAR1 receptor and the CTF of GPR133, we show that acutely induced dissociation of the NTF and thus liberation of the CTF at the plasma membrane increases canonical signaling of GPR133. These findings favor an NTF-CTF dissociation model for activation of GPR133 signaling.

## RESULTS

### Uncleavable GPR133 generates less cAMP signaling relative to WT GPR133

Previous reports suggested canonical signaling by GPR133 is mediated via coupling to G*α*_S_, resulting in an increase of intracellular cAMP (Bohnekamp and Schoneberg, 2011). We independently confirmed that expression of GPR133 in HEK293T cells is associated with robust increase in cAMP levels as detected by a cAMP response element (CRE)-Luciferase reporter, but not other known GPCR signaling pathways (**Figure S1A**). To test whether intramolecular cleavage has implications for canonical signaling, we generated an H543R mutant GPR133 carrying a point mutation at the -2 residue of the GPS cleavage site (Bohnekamp and Schoneberg, 2011) (**Figures 1A-C**). This mutation is known to abolish GPR133 cleavage, but still permits cAMP signaling (Bohnekamp and Schoneberg, 2011; Liebscher et al., 2014). We used homogenous time resolved fluorescence (HTRF) assays to measure the cAMP levels produced by the cleaved WT and uncleaved mutant receptor in HEK293T cells. While overexpression of either receptor variant significantly raised cAMP levels above background, indicating a high baseline signaling activity, overexpression of the uncleavable H543R mutant GPR133 increased cAMP levels only to ∼60% of the cAMP levels obtained with WT-GPR133 (**Figure 1D**). This difference in signaling intensity could not be explained by differences in expression levels of the constructs as assessed by enzyme-linked immunosorbent assays (ELISA) (**Figure 1E**). Similar results were obtained in a patient-derived GBM culture (**Figure 1G-H**). These findings suggest that autoproteolytic cleavage might promote receptor activation.

**Figure 1.**
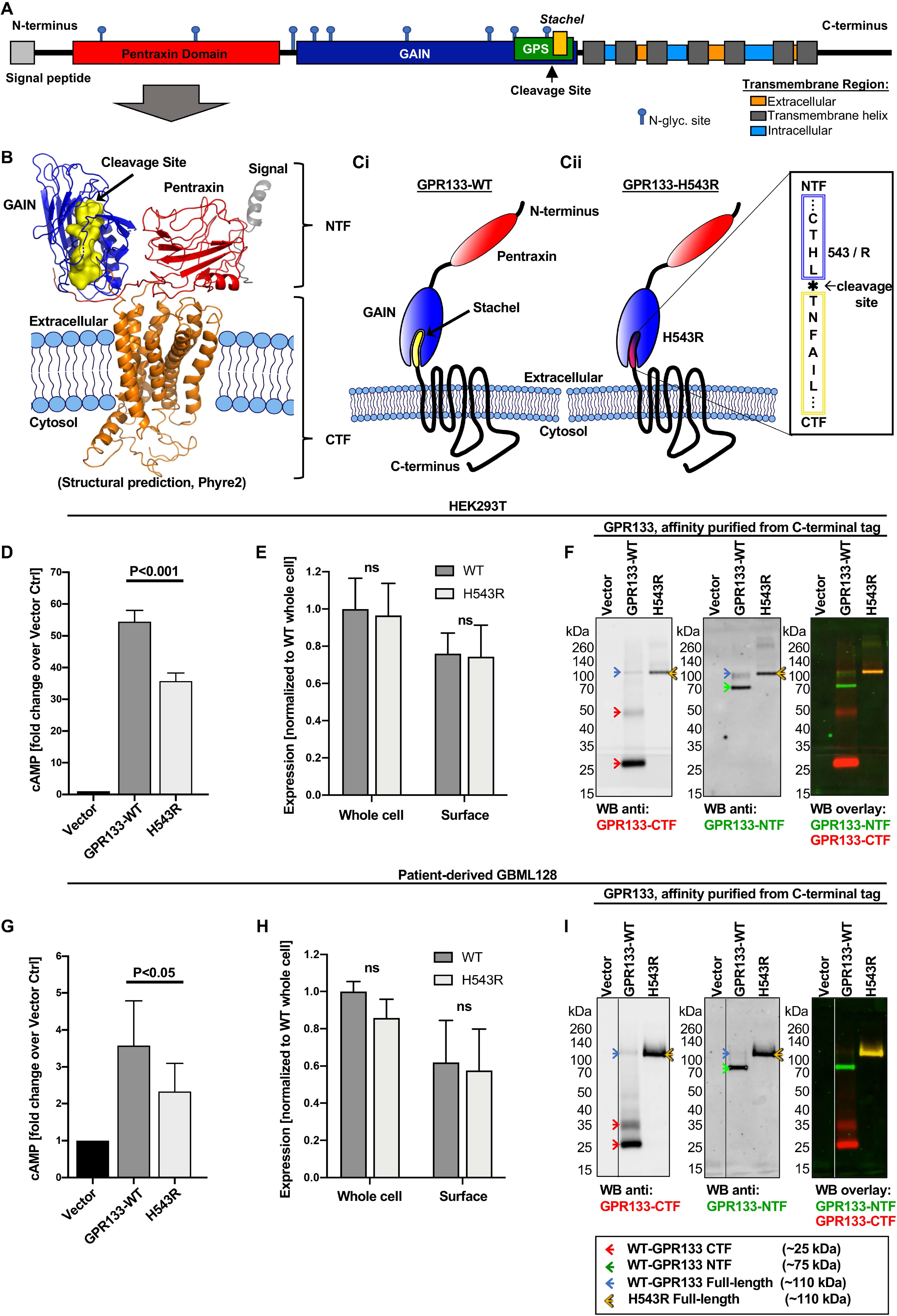
Wild-type GPR133 is cleaved in patient-derived GBM and HEK293T cells and displays higher cAMP signaling relative to uncleaved GPR133. **A)** 2D domain architecture of GPR133 drawn to scale. GAIN, G protein-coupled receptor (GPCR) autoproteolysis inducing domain; GPS, GPCR proteolysis site; CTF, C-terminal fragment; NTF, N-terminal fragment. **B)** 3D protein structure prediction for GPR133 modeled by homology using the Phyre2 web portal (http://www.sbg.bio.ic.ac.uk/~phyre2/html/page.cgi?id=index) (Kelley et al., 2015). The *Stachel* region (yellow surface model) is part of the CTF after cleavage but remains enveloped by the GAIN domain of the NTF (blue ribbon model) in the full-length model. This predicted structure has not been experimentally validated. **C)** Cartoon schematics of wild-type (WT) (**Ci**) and H543R point mutant GPR133 (**Cii**). After cleavage, WT GPR133 is a non-covalently bound heterodimer between the membrane-tethered CTF and the extracellular NTF. The H543R mutation at the -2 residue of the cleavage site prevents intramolecular cleavage and thereby preserves the full-length receptor structure. **D)** Overexpression of WT GPR133 in HEK293T cells results in significantly higher intracellular cAMP levels when compared to the H543R mutant GPR133 as assessed by HTRF assays (cAMP fold change over vector control expressed as mean ± SEM: WT = 54.5 ± 3.6; H543R = 35.8 ± 2.5; ANOVA, F_(2,18)_=115.7, P<0.0001; Tukey’s multiple comparisons adjusted P-value WT vs. H543R: P<0.001; n=7 independent experiments with technical triplicates). **E)** Whole cell- and surface ELISA confirms that expression levels of exogenous WT and H543R mutant GPR133 do not significantly differ from each other in HEK293T cells (Two-way ANOVA, receptor construct F_(1,12)_=2.17, not significant, Whole cell vs Surface F_(1,12)_=0.03, not significant, interaction of factors F_(1,8)_=0.003, not significant; Tukey’s multiple comparisons, all not significant, n=4 independent experiments with technical triplicates). **F)** Multiplexed fluorescent Western blot analysis of GPR133 cleavage products indicates near complete cleavage of WT GPR133 in HEK293T cells. Cells were transfected with an empty vector control, WT, or H543R mutated GPR133. GPR133 fragments were purified from whole cell lysates using a C-terminal TwinStrep affinity tag to reduce non-specific background staining. Western blot membranes were co-stained against the GPR133 CTF (left panel, and red staining in WB overlay) and the GPR133 NTF (middle panel, and green staining in WB overlay). Cleaved GPR133 CTF bands (25 kDa monomer, 48 kDa presumed multimer), cleaved GPR133 NTF bands (75 kDa), WT GPR133 uncleaved bands (∼110 kDa), and the uncleaved H543R mutant GPR133 band (∼110 kDa) are highlighted with red, green, blue, and yellow arrows respectively. A representative blot is depicted. Corresponding whole cell lysate input samples are depicted in **Figure S1B**. **G)** Overexpression of WT GPR133 in patient-derived GBM cells results in significantly higher intracellular cAMP levels when compared to the H543R mutant GPR133 as assessed by HTRF assays (cAMP fold change over vector control expressed as mean ± SEM: WT = 3.58 ± 1.21; H543R = 2.33 ± 0.76; two-tailed ratio paired t-test P-value WT vs. H543R: P<0.05; n=5 independent experiments with technical triplicates). **H)** Whole cell- and surface ELISA confirms that expression levels of exogenous WT and H543R mutant GPR133 do not significantly differ from each other in patient-derived GBM cells (Two-way ANOVA, receptor construct F_(1,8)_=3.88, not significant, Whole cell vs Surface F_(1,8)_=0.30, not significant, interaction of factors F_(1,8)_=0.08, not significant; Tukey’s multiple comparisons, all not significant, n=3 independent experiments with technical triplicates). **I)** WT GPR133 is almost entirely cleaved in patient-derived GBM cells while H543R mutant GPR133 remains uncleaved. GBM cells were lentivirally transduced with an empty vector control, WT, or H543R mutated GPR133, and analyzed as described in Figure 1F. A representative blot is depicted. Corresponding whole cell lysate input samples of this and three additional patient-derived GBM cultures are depicted in **Figure S1C**.

### Wild-type GPR133 is cleaved in patient-derived GBM and HEK293T cells

To assess the extent of intramolecular cleavage of GPR133, we overexpressed WT GPR133 and the H543R point mutant in HEK293T cells and analyzed whole cell lysates by Western blot. Multiplexed fluorescent staining with both a commercial antibody (Sigma, HPA042395) against the GPR133 C-terminal fragment (CTF) and our own previously described monoclonal antibody against the GPR133 N-terminal fragment (NTF) (Bayin et al., 2016; Frenster et al., 2020) detected separate and distinct bands for the CTF (∼25 kDa) and the NTF (∼75 kDa), respectively (**Figures S1B**; red arrows mark the CTF, green arrows mark the NTF). Affinity-purifying the receptor fragments to reduce non-specific background staining confirmed these distinct CTF and NTF bands with increased clarity (**Figures 1F**; red arrows mark the CTF, green arrows mark the NTF). For the WT receptor, we also identified a faint band at 110 kDa (blue arrow), which we hypothesized to represent the uncleaved WT GPR133, as well as a faint band around 48 kDa (red arrow), possibly a dimer of the CTF (**Figures 1F** and **Figure S1B**). In contrast, the H543R mutated GPR133 is detected as a single full-length band (∼110 kDa, yellow arrow), consistent with its cleavage deficiency (**Figure 1F** and **Figure S1B**; yellow arrows). The same cleavage pattern was confirmed in four separate patient-derived GBM cell cultures (affinity-purified receptor fragments in **Figure 1I** and whole cell lysates in **Figure S1C**). It is important to comment on the discrepancies between expected and observed molecular weights (MW) of the uncleaved receptor, NTF, and CTF. The expected MW of the uncleaved receptor without the signal peptide is 93 kDa, while the NTF and CTF are expected at 57 kDa (without the signal peptide) and 36 kDa, respectively. The shifts in observed MW of the uncleaved receptor and NTF are due to glycosylation, as demonstrated later in the manuscript. The shift in the MW of the CTF from 36 kDa (expected) to 25 kDa (observed) is likely explained by increased sodium dodecyl sulfate (SDS) loading on the helical hydrophobic transmembrane segments of the CTF, as previously reported for other transmembrane proteins (Rath et al., 2009).

Overall, these findings suggest that GPR133 is almost entirely cleaved in human GBM and HEK293T cells.

### Intramolecular cleavage of GPR133 is not required for subcellular trafficking to the plasma membrane

To understand mechanisms underlying the increased signaling generated by WT GPR133 compared to the uncleavable mutant receptor, we first analyzed their trafficking to the plasma membrane through the secretory pathway. Using confocal microscopy and indirect immunofluorescent staining under non-permeabilizing conditions, we detected both the WT and the H543R mutated GPR133 at the plasma membrane of cells (**Figure 2Ai-iii** and **Figure S2A**). Similarly, under permeabilizing conditions, the WT and the H543R mutated GPR133 demonstrated analogous staining patterns in both intracellular organelles of the secretory pathway, as well as at the plasma membrane (**Figure 2Bi-iii**).

To confirm these findings biochemically, we used subcellular fractionation and Western blot analysis to separately interrogate three fractions enriched for 1) cytosol, with some endoplasmic reticulum (ER) contamination (Cyto/ER), 2) nucleus, ER, and the Golgi apparatus (Nuc/ER/Golgi), and 3) the plasma membrane (PM). It is noteworthy that while the first two fractions showed enrichment for distinct subcellular compartments/organelles, the plasma membrane fraction was highly specific, as demonstrated by absence of staining for any non-plasma membrane compartment markers by Western blot (**Figure 2E**). Both the cleaved NTF and CTF of the WT receptor, as well as the uncleaved H543R mutant, were prominently detected in the PM fraction (**Figures 2C, D**, red arrowheads), consistent with our microscopy data. Collectively, these findings suggested that intramolecular cleavage of GPR133 is not required for subcellular trafficking to the plasma membrane and that the observed difference in signaling intensities between the cleaved and uncleaved GPR133 variants is not likely to be caused by subcellular trafficking defects.

**Figure 2.**
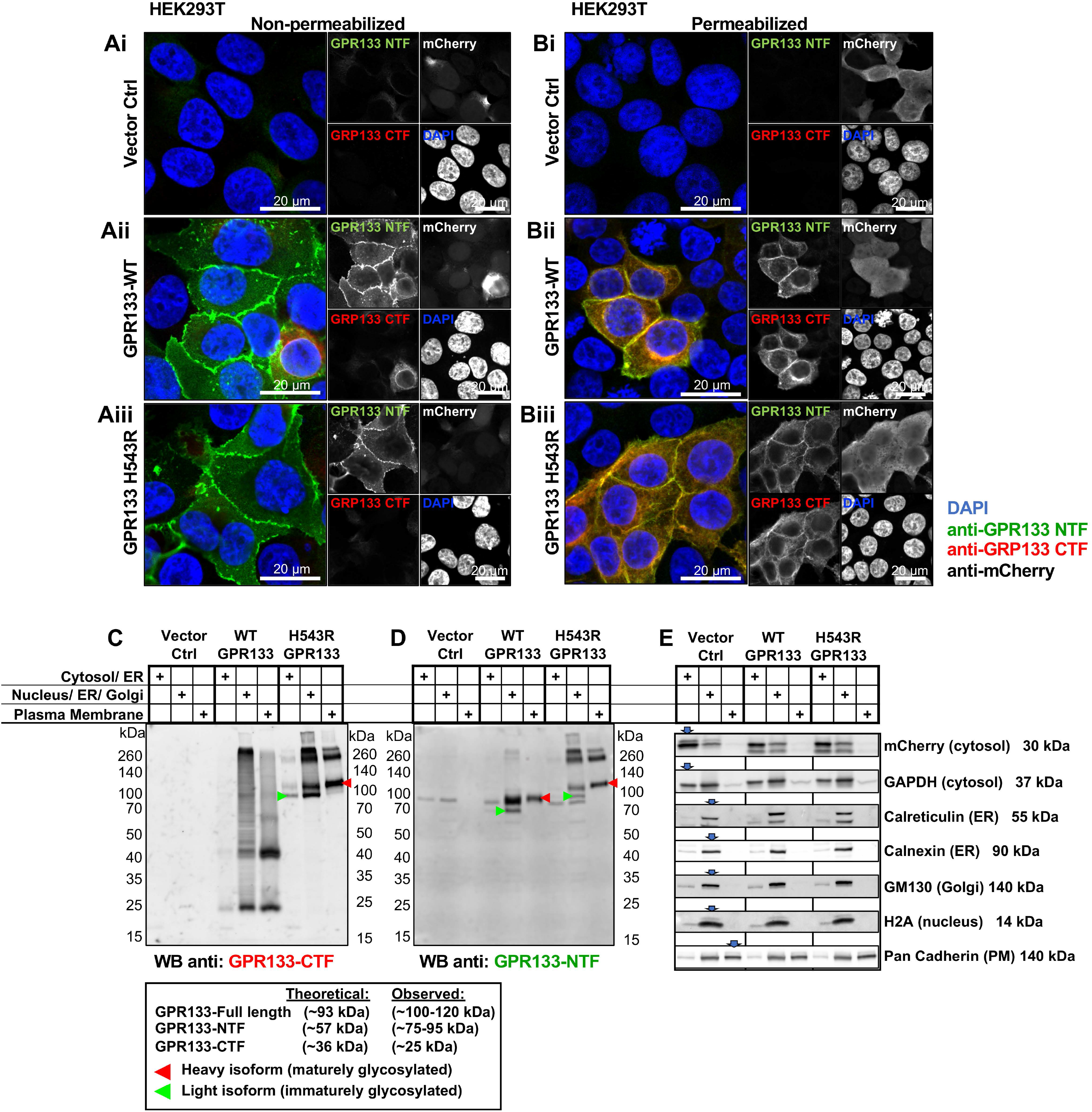
Intramolecular cleavage of GPR133 is not required for subcellular trafficking to the plasma membrane. **A, B)** Representative confocal microscopy micrographs of HEK239T cells overexpressing GPR133 show comparable plasma membrane expression patterns for the WT and H543R mutated GPR133. HEK293T cells were transfected with either an empty vector control (**Ai, Bi**), WT GPR133 (**Aii, Bii**), or H543R mutated GPR133 (**Aiii, Biii**), stained by indirect immunofluorescence under either non-permeabilizing (**A**) or permeabilizing (**B**) conditions. mCherry is co-expressed on all vectors used in this study and is included in the single-channel panels as transfection control (detected by anti-mCherry antibody staining) but is not included in the merged composite panels. Nuclei were counterstained with DAPI. Scale bars, 20 µm. **C, D)** Subcellular fractionation of HEK293T cells expressing an empty vector control (lanes 1-3), WT (lanes 4-6), or H543R mutated GPR133 (lanes 7-9). A representative Western blot stained against the GPR133 CTF (**C**) and the NTF (**D**) is depicted. Both WT and H543R mutated GPR133 are detected in the plasma membrane fractions. Lower molecular weight bands of immaturely glycosylated isoforms are highlighted with green arrowheads (WT-NTF at ∼75 kDa, H543R full-length at ∼100 kDa), higher molecular weight bands of maturely glycosylated isoforms are highlighted with red arrowheads (WT-NTF at ∼95 kDa, H543R full-length at ∼120 kDa). Comparable results were obtained in n=5 independent experiments. The distribution of the receptor fragments across the subcellular fractions is quantified in Figure 4 C. **E)** Subcellular compartment markers validate the specific enrichment of the subcellular fractions as annotated. Panels **C-E** depict the same samples of a representative subcellular fractionation.

### Intramolecular cleavage of GPR133 happens early in the secretory pathway and prior to mature glycosylation

While interrogating the distribution of GPR133 across the subcellular fractions, it became apparent that the cleaved WT GPR133 NTF and the H543R full-length GPR133 undergo a MW shift from lower weight bands in the fractions containing proteins from the early secretory pathway towards higher molecular weight bands in the plasma membrane fraction (**Figure 2C, D**, green and red arrowheads respectively). Since there are nine N-linked glycosylation sites predicted within the NTF (**Figure 1A**), we hypothesized that these observed size shifts are due to different extents of glycosylation as the receptor matures through the secretory pathway. To test this hypothesis, we treated the different subcellular fractions with an enzymatic deglycosylation mix (containing: PNGase F, O-glycosidase, α2-3,6,8,9 neuraminidase A, β1-4 galactosidase S, and β-N-acetylhexosaminidase_f_). Indeed, upon deglycosylation, the different MW isoforms of cleaved WT-NTF and full-length H543R GPR133 shifted to the same predicted MW independent of their subcellular fraction of origin (**Figure 3A**, green, red, and blue arrowheads demarking immaturely glycosylated, maturely glycosylated, and deglycosylated bands, respectively; deglycosylated whole cell lysates are shown in **Figure S3A** for reference), confirming that these bands represent the same protein with different extents of glycosylation.

**Figure 3.**
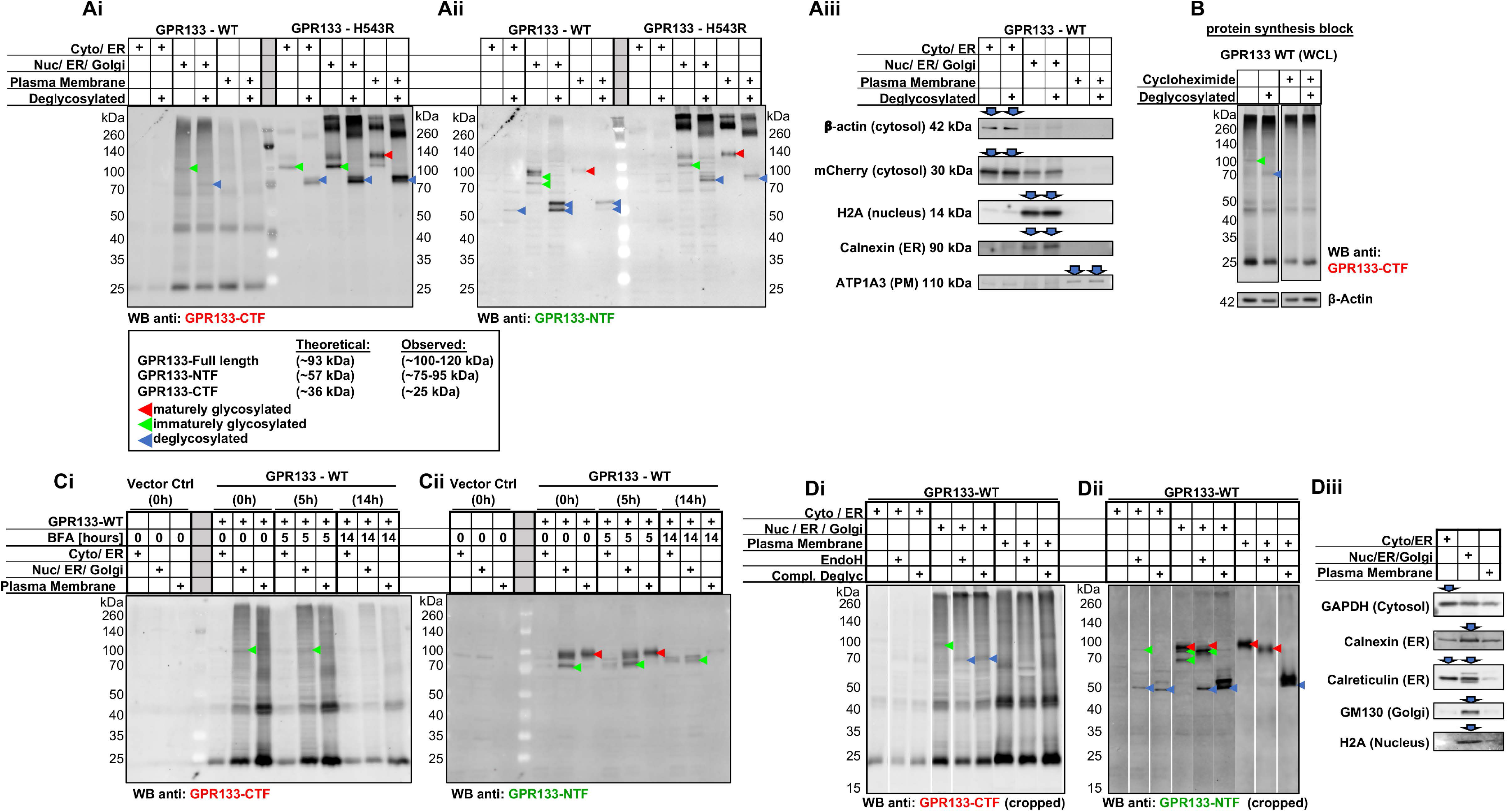
Intramolecular cleavage of GPR133 happens early in the secretory pathway and prior to mature glycosylation. Presumed maturely glycosylated, immaturely glycosylated, and completely deglycosylated forms of GPR133 are marked with red, green, and blue arrowheads respectively, throughout. **A)** Subcellular fractionation of HEK293T cells expressing WT or H543R mutant GPR133 followed by complete enzymatic deglycosylation. GPR133 isoforms were analyzed by Western blot and simultaneously co-stained with an antibody against the GPR133 CTF (**Ai**) and the GPR133 NTF (**Aii**) in separate fluorescent channels. Immaturely and maturely glycosylated isoforms of GPR133 (green and red arrowheads respectively) converge at the same molecular weight upon deglycosylation (blue arrowheads), as observed in both the full-length GPR133 and the cleaved NTF (**Ai-ii**). Note that uncleaved WT GPR133 is detected in the Nuc/ER/Golgi fraction (**Ai**, green and blue arrowheads). Subcellular compartment markers validate enrichment of the respective subcellular fractions (**Aiii**). **B)** Blocking protein synthesis with cycloheximide abolishes the uncleaved WT GPR133 isoform (green and blue arrowheads in untreated sample). HEK293T cells overexpressing WT GPR133 were treated with cycloheximide (280 μg/mL) or vehicle control for 8 hours. Whole cell lysates were deglycosylated and analyzed by Western blot using the GPR133 CTF targeting antibody. β-Actin loading controls from both panels are cropped from the same membrane and exposure. Representative blots from 4 independent experiments are depicted. **C)** Blocking ER-to-Golgi transport with brefeldin A (BFA) does not result in the accumulation of the uncleaved WT GPR133 isoform (**Ci**, green arrowheads). GPR133 overexpressing HEK293T cells were treated with 3 μg/mL BFA as annotated, lysed, and subjected to subcellular fractionation. Confocal microscopy micrographs of GPR133 localization after BFA treatment are depicted in **Figure S3B**. **D)** Lysates of HEK293T cells overexpressing WT GPR133 were subjected to subcellular fractionation followed by treatment with endoglycosidase H (EndoH) or complete enzymatic deglycosylation mix. Note that the uncleaved form of WT GPR133 is sensitive to EndoH treatment (**Di**, green and blue arrowheads), and that the cleaved NTF exists a both immaturely glycosylated EndoH-sensitive, and maturely glycosylated EndoH-insensitive forms (**Dii**). Uncropped membranes are depicted in **Figure S3C**. Subcellular compartment markers in panel **Diii** validate enrichment of the respective subcellular fractions for samples in panels **C** and **D**.

When staining WT GPR133 using the anti-CTF antibody, we detected a band in the Nuc/ER/Golgi fraction shifting from ∼100 kDa to ∼70 kDa upon deglycosylation (**Figure 3Ai**, green and blue arrowheads furthest left). This pattern mimics the immaturely glycosylated uncleaved H543R GPR133, leading to our hypothesis that this band is the low-abundance uncleaved form of WT GPR133 we had previously observed in Western blots from whole cell lysates (**Figure 1F, I**). The faster than expected mobility of this uncleaved deglycosylated form likely results from increased SDS binding on the helical transmembrane segments of the receptor (Rath et al., 2009). To test whether this uncleaved and immaturely glycosylated form of WT GPR133 is a stable form of the receptor or a transient state during receptor maturation, we blocked protein synthesis with cycloheximide. Protein synthesis block indeed abolished this uncleaved form of GPR133, supporting the hypothesis that the uncleaved WT GPR133 is a short-lived transition state (**Figure 3B**, green and blue arrowheads).

To determine whether this short-lived full-length WT GPR133 is cleaved directly after synthesis in the ER, or later in the Golgi apparatus, we treated cells with Brefeldin A (BFA), which interrupts ER-to-Golgi transport. The effectiveness of BFA to prevent transport along the secretory pathway was confirmed by confocal microscopy (**Figure S3B**). Brefeldin A treatment did not lead to an accumulation of the uncleaved WT GPR133 isoform (**Figure 3Ci**, green arrowheads), suggesting cleavage happens immediately after protein synthesis. However, BFA did result in the elimination of maturely glycosylated NTF in the Nuc/ER/Golgi fraction 14 hours after its addition to the medium (**Figure 3Cii**, red arrowheads).

Lastly, we treated the subcellular fractions with endoglycosidase H (EndoH), which removes the immature high mannose glycosylation of proteins within the ER but not mature glycosylation of proteins that have reached the Golgi apparatus. Indeed, we observed that the uncleaved WT GPR133 band was sensitive to EndoH-mediated deglycosylation without additional deglycosylation effect conferred by a complete deglycosylation mix containing PNGase, suggesting it represents an immaturely glycosylated protein localizing to the ER (**Figure 3D** and **Figure S3C**).

These data suggest a model in which newly synthesized WT GPR133 carrying immature glycosylation gets intramolecularly cleaved within the ER prior to trafficking to the Golgi apparatus, where it acquires mature glycosylation. WT GPR133 reaches the plasma membrane as a fully cleaved and maturely glycosylated protein.

### The GPR133 NTF dissociates from the CTF at the plasma membrane

To investigate whether the cleaved GPR133 CTF and NTF remain non-covalently bound to each other, we created GPR133 constructs carrying either C-terminal or N-terminal TwinStrep-tags for affinity purification (**Figure 4A**). When purifying the cleaved receptor from whole cell lysates, identical stoichiometries of CTF and NTF were eluted, independent of whether they are purified using the N-terminal or C-terminal affinity tag (**Figure 4B**).

**Figure 4.**
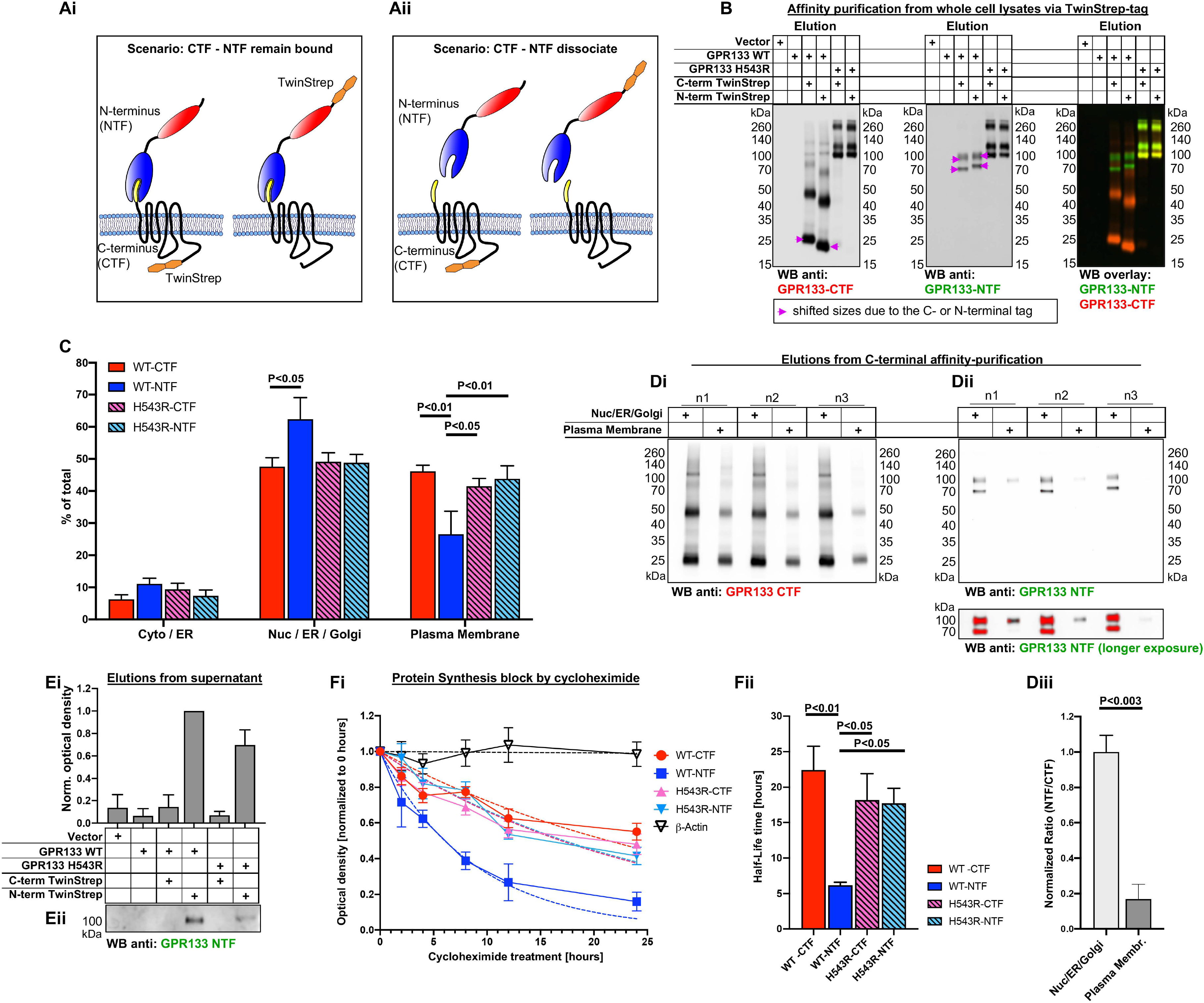
The GPR133 NTF dissociates from the CTF at the plasma membrane. **A)** Schematic of C-terminally and N-terminally tagged GPR133 constructs and experimental rationale. Conceptually, if the CTF and NTF remain bound to each other, receptor constructs with C-terminal or N-terminal affinity tags should purify the CTF and NTF at a one-to-one ratio (**Ai**). In the event of dissociation, the location of the affinity tag would bias the ratio of CTF or NTF in the elution (**Aii**). **B)** Western blot analysis of GPR133 constructs affinity-purified from whole cell lysates. On a whole cell lysate level, both the C-terminally and N-terminally tagged GPR133 constructs copurify the CTF and NTF at similar stoichiometry, indicating the CTF and NTF to be mostly bound as heterodimer. **C)** Quantified distribution of GPR133 fragments across different subcellular fractions detects relatively less WT GPR133 NTF than CTF at the plasma membrane. HEK293T cells overexpressing either WT or H543R mutated GPR133 were subjected to subcellular fractionation, followed by Western blotting. The subcellular distribution of each receptor fragment as detected by the NTF- or CTF-targeting antibody was quantified by densitometry. Results are depicted as mean ± SEM (Two-way ANOVA, receptor fragment F_(3,48)_=3.41×10^-8^, P>0.99, subcellular fraction F_(2,48)_=153.4, P<0.0001, interaction of factors F_(6,48)_=5.02, P<0.001; Tukey’s multiple comparisons Nuc/ER/Golgi WT-CTF vs. WT-NTF P<0.05; Plasma Membrane WT-CTF vs. WT-NTF P<0.003, WT-NTF vs. H543R-CTF P<0.03, WT-NTF vs. H543R-NTF P<0.01, n=5 independent experiments). A representative Western blot membrane of this distribution is depicted as part of Figure 2 C,D. **D)** Tagged GPR133 CTF copurifies less NTF from the plasma membrane fraction than from the secretory pathway. HEK293T cells overexpressing C-terminally tagged WT GPR133 were subjected to subcellular fractionation, followed by affinity purification of the GPR133 CTF. Elutions from the Nuc/ER/Golgi and the plasma membrane fractions were analyzed by Western blot (**Di-ii**) and intensities were quantified by densitometry. **Diii)** Summary of quantified NTF/CTF ratios from Di-ii depicted as mean ± SEM. (Nuc/ER/Golgi: 1.00±0.09; plasma membrane: 0.17±0.08; unpaired t-test P<0.003; n=3 independent experiments). Upper and lower panels of **Dii** depict the same membrane at different exposures. Red indicates signal saturation. **E)** Soluble NTF is detected in cell culture supernatants. Supernatants from HEK293T cells overexpressing different variants of GPR133 were collected and precleared by centrifugation. Tagged receptor fragments were affinity-purified and analyzed by Western blot (**Eii**). The quantified relative densities are depicted as mean ± SEM (**Ei**). No staining for the receptor CTF was detected in any condition. Corresponding whole cell lysates of the overexpressing cells, are depicted in panel **B**. Full-size Western blot membranes for the CTF and NTF staining and deglycosylation of the soluble NTF fragment are depicted in **Figure S4A-B**. **F)** GPR133 protein half-life and decay curves after protein synthesis block with cycloheximide. HEK293T cells overexpressing either WT or H543R mutated GPR133 were treated with 280μg/mL cycloheximide and harvested for analysis at different time points. Whole cell lysates were analyzed by Western blot, and relative abundance of receptor fragments as detected with the CTF or NTF-targeting antibodies were quantified. **Fi)** Protein amounts normalized to the beginning of cycloheximide chase are plotted as a function of time. Dotted lines are modeled one-phase decay curves used to calculate half-life times. **Fii)** Protein half-life times of GPR133 fragments depicted as mean ± SEM. The WT-NTF has a significantly shorter half-life in whole cell lysates than the CTF or the uncleaved receptor (ANOVA F_(3,12)_=6.47, P<0.008; Tukey’s multiple comparisons: WT-CTF vs WT-NTF: P<0.006; WT-NTF vs H543R-CTF: P<0.05; WT-NTF vs H543R-NTF: P<0.05; n=4 independent experiments). Representative Western blots of the cycloheximide time-course stained against the CTF and NTF of GPR133 are depicted in **Figure S4Ci-iv**.

While this finding suggested that the NTF and CTF remain non-covalently associated at the whole cell level, we proceeded to dissect whether this association varied depending on the location of GPR133 along the secretory pathway. Towards this, we quantified the relative distribution of each receptor fragment (WT-CTF, WT-NTF, H543R full-length as detected with the CTF antibody, H543R full-length as detected with the NTF antibody) across the three subcellular fractions mentioned above (**Figure 4C**). The subcellular distribution did not differ between the cleaved WT-CTF and the uncleaved H543R mutant using either the C-terminal or N-terminal antibody, supporting the previous observation that cleavage does not affect the trafficking of the receptor’s transmembrane segment. However, the cleaved WT-NTF was significantly underrepresented at the plasma membrane when compared to the WT-CTF or the H543R uncleaved receptor (**Figure 4C**). We note that, while the WT-NTF appears to be overrepresented in the Nuc/ER/Golgi fraction compared to the WT-CTF, this is likely a mathematical artifact arising from reduced representation of the WT-NTF in the plasma membrane fraction. This finding suggested that the cleaved NTF either traffics within the cells independently of the CTF, or more likely, that the NTF dissociates from the CTF at the plasma membrane and is thus less abundant in that fraction.

To test these two models, we repeated the purification of the receptor using its C-terminal affinity tag and compared the Nuc/ER/Golgi fraction against the plasma membrane fraction as input. We detected a significantly lower NTF-to-CTF ratio at the plasma membrane when compared to the Nuc/ER/Golgi fraction (**Figure 4Di-iii**). Furthermore, we were able to purify and detect the soluble NTF from precleared cell culture supernatants, while not detecting any associated C-terminal fragments (**Figure 4E** and **Figure S4A-B**). These findings suggested that, while the cleaved NTF and CTF are non-covalently bound in the secretory pathway, they do partially dissociate at the plasma membrane.

To gather additional evidence for such NTF-CTF dissociation at the plasma membrane, we assayed protein decay of the two fragments after blocking protein synthesis with cycloheximide in HEK293T cells overexpressing either the cleaved WT or uncleaved H543R mutant receptor (**Figure 4 Fi-Fii** and **Figure S4C**). Whole cell lysates of different cycloheximide chase time points were analyzed by Western blot using our CTF and NTF targeting antibodies and intensities were plotted as a function of time. No significant difference was detected between the decay curves and half-life times of uncleaved H543R GPR133 and the WT-CTF, indicating that the above described signaling intensity differences were not caused by differences in protein stability. However, the cleaved WT GPR133 NTF decayed at a significantly faster rate than the WT-CTF. While this data could be interpreted as accelerated degradation of the NTF compared to the CTF, it more likely suggests loss of NTF from whole cell lysates due to its dissociation and diffusion into the supernatant. This latter interpretation of soluble NTF is supported by the aforementioned fact that NTF was detected in precleared cell culture supernatants (**Figure 4E**), and the fact that we did not detect exogenous NTF on the surface of cells adjacent to GPR133-overexpressing cells (**Figure S4D, E**). Therefore, this difference in decay curves offers additional support to the hypothesis that the NTF and CTF time-dependently dissociate at the plasma membrane.

### NTF dissociation at the plasma membrane increases canonical signaling of a hybrid PAR1-GPR133 receptor

The data above suggest that the NTF is shed at the plasma membrane, which may explain the elevated signaling of cleaved WT GPR133 relative to the uncleavable receptor. To directly test whether NTF dissociation is the mechanism responsible for increased signaling, we constructed a fusion protein between the N-terminus of protease-activated receptor 1 (hPAR1) and the CTF of GPR133, following the design of Mathiasen and colleagues (Mathiasen et al., 2020). In this fusion protein, the PAR1 NTF serves as a proxy for the NTF of GPR133 by capping the GPR133 CTF during maturation, while also adding a thrombin recognition site for enzymatically inducible cleavage and release of the NTF at the plasma membrane (**Figures 5A** and **Figure S5A**). At the cleavage side of this fusion protein, the residues “SF” of hPAR1 replace the residues “TNF” of GPR133’s *Stachel* region, thus labeled “PAR1-GPR133 *Δ*TN” (fusion construct and tested alternative designs detailed in **Figure S5**). Using patient-derived GBM cells, we indeed observed that thrombin-mediated cleavage of the fusion protein resulted in a significant increase in intracellular cAMP levels in a concentration-dependent manner (**Figure 5B** and **Figure S5Bi-ii**). WT GPR133 lacking the thrombin recognition site did not respond to thrombin treatment, supporting the specificity of this effect (**Figure 5B** and **Figure S5Bi-ii**). Similar results were obtained in HEK293T cells (**Figure 5D** and **Figure S5Di-Dii**). Using cell surface enzyme-linked immunosorbent assays (ELISAs) in HEK293T cells, we demonstrated that this exposure to thrombin indeed led to the dissociation of the PAR1-NTF, with a concentration-dependence that paralleled our signaling data (**Figure 5C** and **Figure S5Ci-ii**). These findings support the hypothesis that NTF dissociation promotes full activation of GPR133.

**Figure 5.**
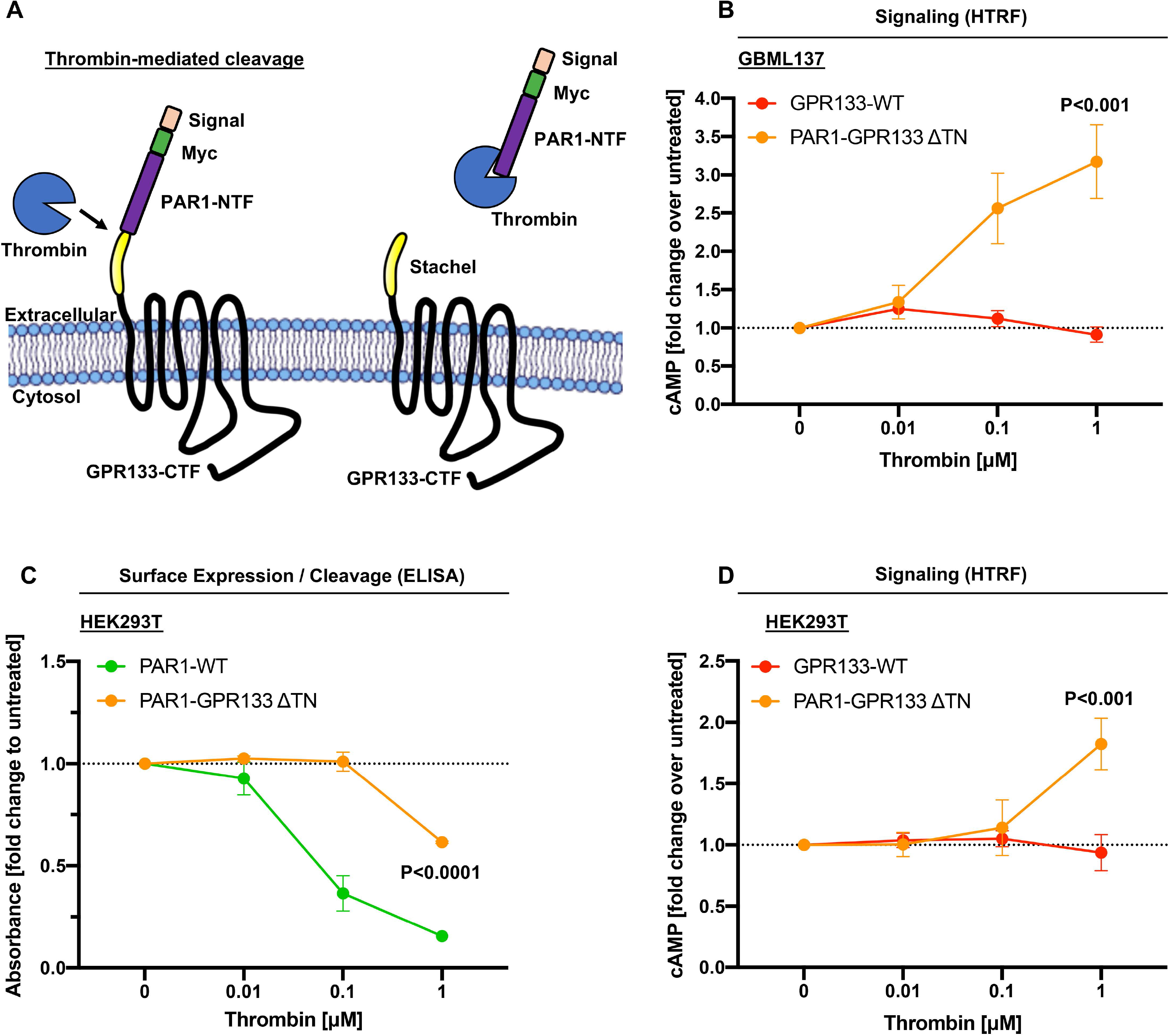
NTF shedding at the plasma membrane increases canonical signaling of a hybrid PAR1-GPR133 receptor. **A)** Cartoon schematic of thrombin-mediated cleavage of the PAR1-GPR133 fusion protein. Myc-tagged human PAR1 NTF including its thrombin recognition site and subsequent two amino acids (SF) was fused to the GPR133 CTF. The last two amino acids of the PAR1 NTF replaced the first three residues of the GPR133 *Stachel* sequence (TNFàSF, called “ΔTN”). Successful thrombin-mediated cleavage of this construct was assessed by loss of the N-terminal Myc-tag. Detailed information on this and additional fusion constructs is shown in **Figure S5**. **B)** Patient-derived GBM cultures overexpressing either WT GPR133 or the PAR1-GPR133-ΔTN fusion were exposed to varying concentrations of thrombin, and intracellular cAMP levels were assessed via HTRF assays. Data is depicted as mean ± SEM normalized to the untreated condition. Thrombin-mediated cleavage of the NTF significantly increased canonical GPR133 signaling in the PAR1-GPR133-ΔTN fusion, but not WT GPR133 lacking the thrombin recognition site (GBML137: Two-way ANOVA, GPR133 constructs F_(1,16)_=27.73, P<0.0001, thrombin F_(3,16)_=7.17, P<0.003, interaction of factors F_(3,16)_=9.27, P<0.001; Tukey’s multiple comparisons, 1 µM thrombin, GPR133 WT vs PAR1-GPR133-ΔTNF: P<0.001, n=3 independent experiments with technical triplicates). **C)** Surface ELISA detects the thrombin-mediated cleavage of the Myc-tagged PAR1-NTF in both the full-length PAR1 (positive control) and the PAR1-GPR133-ΔTNF fusion protein in a concentration-dependent manner in HEK293T cells (Tukey’s multiple comparisons, PAR1-GPR133-ΔTNF fusion 1 µM thrombin vs untreated: P<0.0001, n=3 independent experiments with technical triplicates). **D)** HEK293T cells overexpressing either WT GPR133 or the PAR1-GPR133-ΔTN fusion were exposed to varying concentrations of thrombin, and intracellular cAMP levels were assessed via HTRF assays. Data is depicted as mean ± SEM normalized to the untreated condition. Thrombin-mediated cleavage of the NTF significantly increased canonical GPR133 signaling in the PAR1-GPR133-ΔTN fusion, but not WT GPR133 lacking the thrombin recognition site (Two-way ANOVA, GPR133 constructs F_(1,16)_=6.61, P<0.03, thrombin F_(3,16)_=3.67, P<0.04, interaction of factors F_(3,16)_=5.68, P<0.008; Tukey’s multiple comparisons, 1 µM thrombin, GPR133 WT vs PAR1-GPR133-ΔTNF: P<0.004, n=3 independent experiments with technical triplicates). Absolute values and additional variations of the PAR1-GPR133 fusion constructs are shown in **Figure S5**.

## DISCUSSION

Classically, GPCRs are thought to exist in an equilibrium between an active and inactive state. Upon encountering a stimulus, such as ligand binding, this equilibrium shifts towards the “on” state by stabilizing the receptor in a certain conformation (Lefkowitz et al., 1993). For adhesion GPCRs, such a shift in the equilibrium is hypothesized to be mediated by a tethered internal agonist, the *Stachel* sequence, which resides in the most N-terminal region of the CTF after cleavage (Liebscher and Schoneberg, 2016; Maser and Calvet, 2020). There are two possible modes for how the *Stachel* sequence might exert its agonistic effect on adhesion GPCR signaling. Either a conformational change within the extracellular region containing the *Stachel* is sufficient for receptor activation, as would be expected from a classical GPCR; or dissociation of the NTF from the CTF causes unmasking of the *Stachel* sequence, which in turn causes receptor activation. The facts that uncleavable mutant adhesion GPCRs in some cases phenocopy the canonical signaling of their wild-type counterparts, and that several adhesion GPCRs are not cleaved, argue for the former model (Kishore et al., 2016; Promel et al., 2012; Wilde et al., 2016). However, examples of cleavage-deficient mutants manifesting reduced signaling capacity (Hsiao et al., 2011; Hsiao et al., 2015; Huang et al., 2012; Zhu et al., 2019), reports of soluble NTFs of adhesion GPCRs detected *in vitro* and *in vivo* (Cork et al., 2012; Huang et al., 2012; Kaur et al., 2005; Vallon and Essler, 2006), and examples of deletion mutants mimicking CTFs that demonstrate increased signaling relative to their wild-type counterparts (Kishore et al., 2016; Liebscher et al., 2014; Paavola et al., 2011), argue in favor of the latter model. In the case of GPR133, where the uncleavable H543R mutant demonstrates 60% of the basal activity of the wild-type cleaved receptor, it is possible that the *Stachel* sequence is already prebound in the agonist binding site of the CTF and cleavage allows full isomerization to the active state. Such a model would be consistent with isomerization properties of other GPCRs with tethered agonists (Bruser et al., 2016; Schoneberg et al., 2016; Schulze et al., 2020).

Our study provides evidence that intramolecular cleavage of GPR133 further increases receptor activity via dissociation of the cleaved N-terminus at the plasma membrane (**Figure 6**). The survey of both patient-derived GBM cells and HEK293T cells indicated that GPR133 is almost entirely cleaved before the receptor reaches the plasma membrane. While Western blot analysis of whole cell lysates did detect a small amount of uncleaved GPR133, we believe that this form of GPR133 represents a transient state of the newly synthesized receptor in the ER. This view is supported by the observations that the uncleaved receptor 1) did not appear in the subcellular fraction representing the plasma membrane (**Figure 2C** and **Figure 3A, C-D**; 2) was sensitive to deglycosylation by EndoH, without additional deglycosylation effect conferred by PNGase (**Figure 3D**); 3) did not accumulate upon BFA treatment (**Figure 3C**); and 4) was abolished by blocking protein synthesis with cycloheximide (**Figure 3B**). Our observations are in agreement with previous reports on the adhesion GPCRs CIRL (ADGRL1) and GPR116 (ADGRF5) (Abe et al., 2002; Krasnoperov et al., 2002).

**Figure 6.**
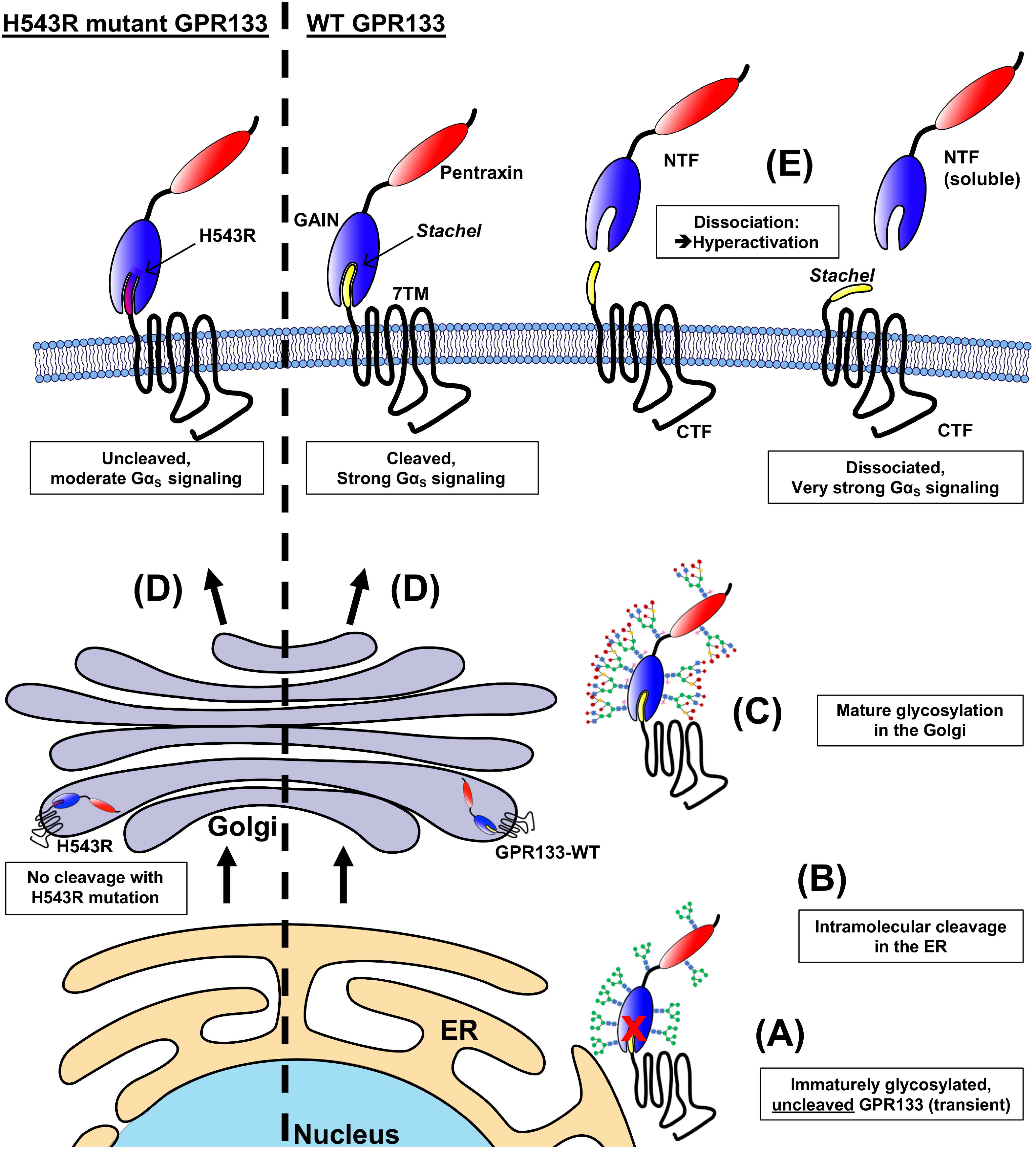
Proposed model of GPR133’s molecular life cycle. **(A)** Newly synthesized GPR133 in the endoplasmic reticulum (ER) is immaturely glycosylated and uncleaved. **(B)** Wild-type (WT) GPR133 is intramolecularly cleaved in the ER prior to mature glycosylation. **(C)** GPR133 in the Golgi apparatus gains mature glycosylation on its NTF resulting in a large MW shift by Western blot analysis (Figures 3 and S3). The cleaved CTF and NTF remain non-covalently bound to each other as a heterodimer and copurify at a one-to-one ratio. **(D)** Fully cleaved WT GPR133 and uncleaved H543R mutated GPR133 indistinguishably traffic to the plasma membrane in a maturely glycosylated state. **(E)** The GPR133 NTF dissociates from the CTF resulting in its relatively lower abundance at the plasma membrane and its consequent detection in the supernatant. Acute liberation of the CTF and exposure of the *Stachel* increases canonical signaling of GPR133.

While the intramolecular cleavage of GPR133 takes place early in the secretory pathway, it is not required for glycosylation of the N-terminus and subcellular trafficking of GPR133 to the plasma membrane. Previous literature on this subject has remained controversial, as it contains examples of both appropriate and arrested trafficking of cleavage-deficient point mutant adhesion GPCRs (Arac et al., 2012; Huang et al., 2012; Kishore et al., 2016; Krasnoperov et al., 2002; Moriguchi et al., 2004; Okajima et al., 2010; Volynski et al., 2004). This controversy raises the question whether the requirement of cleavage for correct subcellular trafficking is truly as receptor-specific as it appears from the literature, or whether misfolding due to the introduced mutations at the cleavage site is responsible for the lack of plasma membrane localization in some of the adhesion GPCRs.

Our biochemical analysis indicated that the cleaved NTF and CTF of GPR133 remain non-covalently bound during receptor trafficking, until the NTF dissociates at the plasma membrane. We propose that this dissociation underlies the higher signaling capacity of cleaved GPR133 relative to the uncleavable H543R mutant receptor. The conclusion that the dissociation occurs at the plasma membrane is supported by the following observations: 1) while on a whole cell lysate level the CTF and NTF appeared to co-purify at similar stoichiometries, suggesting non-covalent association (**Figure 4B**), CTF from the plasma membrane fraction was significantly less associated with NTF when compared to CTF from the ER fraction (**Figure 4Di-iii**); 2) the NTF displayed a faster decay curve than CTF in whole cell lysates after protein synthesis block, indicating that CTF and NTF are capable of behaving as distinct entities (**Figure 4Fi-ii**); 3) we detected NTF in the cell culture supernatant, consistent with NTF dissociation from the CTF at the plasma membrane (**Figure 4Ei-ii**). To our knowledge, this is the first experimental evidence of such NTF-CTF dissociation for GPR133.

To demonstrate that cleavage followed by NTF dissociation is not only correlated to increased canonical signaling but that these two aspects are causally related, we used a recently published model for experimentally controlled shedding of the NTF (Mathiasen et al., 2020). We generated a fusion protein between the protease-activated receptor 1 (hPAR1) N-terminus and GPR133’s CTF, and administered thrombin to induce the enzymatic cleavage of the PAR1 NTF and acute dissociation from the CTF. We showed that this acute thrombin-induced NTF shedding at the plasma membrane increases the intensity of canonical signaling of the fusion protein, while not affecting WT GPR133 lacking the thrombin recognition site.

Having demonstrated that GPR133 is intramolecularly cleaved and that the dissociation of the NTF promotes receptor activation and canonical signaling, our future efforts will focus on identifying mechanisms underlying NTF-CTF dissociation at the plasma membrane. One hypothesized mechanism of dissociation is that ligand binding confers mechanical shear stress onto the NTF, thereby pulling it off the CTF (Petersen et al., 2015; Scholz et al., 2016; Stoveken et al., 2015). This model assumes that the strength of the NTF-ligand interaction exceeds that of the NTF-CTF association. An alternative mechanism may involve conformational shifts at the GAIN domain following ligand-NTF binding, leading to weakened NTF-CTF association and shedding, without the need for strong mechanical forces. More likely, a combination of the two scenarios might take place and thus increase the specificity of an activating signal. Other physicochemical parameters may also regulate this dissociation, including pH, stiffness of the extracellular matrix or receptor multimerization.

In conclusion, we demonstrate that dissociation of the cleaved NTF at the plasma membrane increases GPR133 activity and canonical signaling. The fact that the uncleavable GPR133 point mutant is also capable of signaling, albeit to a lesser extent relative to the wild-type receptor, suggests that the steady-state equilibrium allows for a moderate amount of baseline signaling independent of additional stimuli, possibly due to the endogenous *Stachel* sequence being internally prebound in a configuration permissive for signaling. Alternatively, partial activation of the uncleaved receptor may occur through a combination of ligands, mechanical forces or other stimuli that intermittently shift the receptor equilibrium to the “on” state. The possibility of extracellular ligands reversibly activating canonical signaling in the absence of intramolecular cleavage has previously been reported for the adhesion GPCR GPR56 (ADGRG1) (Salzman et al., 2017). However, for the WT GPR133, we postulate that dissociation of the NTF irreversibly activates the CTF to the “on” state until the receptor is internalized, degraded or desensitized. Our findings have implications for basic adhesion GPCR biology, but also help elucidate the function of GPR133 in GBM.

## ACKNOWLEDGEMENTS

We thank the Microscopy core facility at NYU Grossman School of Medicine for assistance with confocal microscopy imaging. The facility is supported by the NYU Perlmutter Cancer Center Support Grant P30CA016087. JDF was supported by a NYSTEM Stem Cell Biology training grant to NYU Grossman School of Medicine (#C322560GG). GS was supported by a DFG postdoctoral fellowship (STE 2843/1-1). NRB was supported by a T32 Cell Biology training grant (T32GM136542) to NYU Grossman School of Medicine. IL and TS were supported by DFG FOR2149 (project numbers 266022790 P4 to TS and P5 to IL), CRC1052 (project number 209933838 B6) and CRC1423 (project number 421152132). DGP was supported by NIH/NINDS R01 NS102665, NYSTEM (NY State Stem Cell Science) IIRP C32595GG, NIH/NIBIB R01 EB028774 (to Dr. Steven Baete at NYU Grossman School of Medicine), NYU Grossman School of Medicine, and DFG (German Research Foundation) FOR2149 as Mercator fellow.

## AUTHOR CONTRIBUTIONS (written in CRediT format)

Conceptualization, J.D.F. and D.G.P.; Investigation, J.D.F.; Methodology, J.D.F. and G.S.; Formal Analysis, J.D.F. and D.G.P.; Resources, D.G.P., N.S., I.L., and T.S.; Writing - Original Draft, J.D.F. and D.G.P.; Writing - Review & Editing, J.D.F., G.S., N.R.B., D.B., J.W., B.K., C.W., N.S., G.R.W., I.L., T.S., and D.G.P.; Visualization, J.D.F.; Funding Acquisition, J.D.F., G.S., N.R.B., I.L., T.S., D.G.P.; Supervision, D.G.P.

## STAR Methods

### Methods Details

#### Cell culture

Patient-derived glioblastoma (GBM) cultures were established and maintained as we previously described (Bayin et al., 2016; Bayin et al., 2017; Frenster and Placantonakis, 2018). In brief, fresh operative specimens were obtained from patients undergoing surgery for resection of GBM after informed consent (NYU IRB study 12-01130). Specimens were mechanically minced using surgical blades followed by enzymatic dissociation using Accutase (Innovative Cell Technologies, Cat# AT104). Cells were either long-term maintained in spheroid suspension cultures on untreated cell culture dishes, or grown as attached cultures on dishes pretreated with poly-L-ornithine (Sigma, Cat# P4957) and laminin (Thermo Fisher, Cat# 23017015). The glioblastoma growth medium consisted of Neurobasal medium (Gibco, Cat# 21103049) supplemented with N2 (Gibco, Cat# 17-502-049), B27 (Gibco, Cat# 12587010), non-essential amino acids (Gibco, Cat# 11140050), GlutaMax (Gibco, Cat# 35050061), and was additionally supplemented with 20 ng/mL recombinant basic Fibroblast Growth Factor (bFGF; R&D, Cat# 233-FB-01M) and 20 ng/mL Epidermal Growth Factor (EGF; R&D, Cat# 236-EG-01M) every other day. Parental tumors of these patient-derived cultures underwent mutational and copy number variation (CNV) profiling (**Table S1**) using a focused next-generation sequencing panel of 50 genes (NYU Oncomine focus assay; **Table S2**) (Hovelson et al., 2015; Mehrotra et al., 2017) on an Ion Torrent S5 instrument. All tumors had a wild-type isocitrate dehydrogenase (IDH) background.

HEK293T (Takara, Cat# 632180) cells were cultured in DMEM (Gibco, Cat# 11965-118) supplemented with 10% fetal bovine serum (Peak Serum, Cat# PS-FB2) and sodium pyruvate (Gibco, Cat# 11360070).

All cells were cultured in humidified cell culture incubators at 37°C balanced with 5% CO_2_. Patient-derived GBM cells were cultured at 4% O_2_, while HEK293T cells were cultured at 21% O_2_.

#### HTRF signaling assays

HEK293T or patient-derived GBM cells were transfected with overexpression plasmids of interest using Lipofectamine 2000 (Invitrogen, Cat# 11668-019) or Lipofectamine 2000 Stem reagent (Thermo Fisher, Cat# STEM00008), respectively, according to the manufacturer’s protocol. 24 hours after transfection, cells were reseeded onto 96-well plates pretreated with poly-L-ornithine (Sigma, Cat# P4957) and laminin (Thermo Fisher, Cat# 23017015) at a density of 75000 cells per well. Two days after transfection, the medium was exchanged for 50 μL of fresh medium with 1 mM 3-isobutyl-1-methylxanthine (IBMX) (Sigma-Aldrich, Cat# I7018-100MG), and cells were incubated at 37°C for an additional 30 minutes. For the thrombin-mediated cleavage experiments, thrombin (Sigma-Aldrich, Cat# T9326-150UN) was added into the medium mix as part of this 30-minute incubation. Cells were lysed and cAMP levels were measured using the cAMP Gs dynamic kit (CisBio, Cat# 62AM4PEC) on the FlexStation 3 (Molecular Devices) according to the manufacturer’s protocol.

#### Enzyme-linked immunosorbent assay (ELISA)

Cells were transfected and reseeded as described for HTRF signaling assays. Forty-eight hours after transfection (and after 30 minutes of thrombin exposure if applicable), cells were washed with HBSS +Ca^2+^/+Mg^2+^ and fixed with 4% paraformaldehyde (PFA, Sigma-Aldrich, Cat# P6148) for 20 minutes at room temperature. For whole cell ELISA under permeabilizing conditions, all following steps were conducted in the presence of 0.1% Triton X-100; for surface ELISA under non-permeabilizing conditions no detergent was added at any time. Cells were blocked in HEK293T media containing 10% FBS for 1 hour at room temperature. Cells were incubated with primary antibodies diluted in HEK293T medium containing 10% FBS at concentrations indicated in **Table S3** for 1 hour at room temperature. After 3 washing steps with PBS, cells were incubated with horseradish peroxidase-conjugated secondary antibodies diluted 1:1,000 in HEK293T medium containing 10% FBS for 1 hour at room temperature. After additional 3 thorough washes with PBS, cells were overlayed with TMB-stabilized chromogen for 20 minutes (Thermo Fisher, Cat# SB02) followed by an equal volume of acidic stop solution (Thermo Fisher, Cat# SB04)

#### Chemical drug treatments

Cycloheximide (Sigma-Aldrich, Cat# 239765) was administered to cell culture medium at a final concentration of 280 μg/mL to achieve protein synthesis block. Brefeldin A (BFA) (Invitrogen, Cat# 00-4506-51) was administered to cell culture medium at a final concentration of 3 μg/mL to block ER-to-Golgi transport. 3-isobutyl-1-methylxanthine (IBMX) (Sigma-Aldrich, Cat# I7018) was administered to cell culture medium at a final concentration of 1 mM to block phosphodiesterases during cAMP measurements.

#### Luciferase vector signaling assays

HEK293T cells were co-transfected with GPR133 overexpression plasmids and either one of the following luciferase signaling-reporter plasmids: CRE-Luciferase (Promega, Cat# E8471), SRE-Luciferase (Promega, Cat# E1340), SRF-RE-Luciferase (Promega, Cat# E1350), NFAT-RE-Luciferase (Promega, Cat# E8481), NF*κ*B-RE-Luciferase (Promega, Cat# E8491). 24 hours after transfections, cells were reseeded in black 96-well plates at a density of 75,000 cells per well. 48 hours after transfection cells were lysed and luciferase activity was detected using the Bright-Glo Luciferase assay system (CisBio, Cat# E2650) and a BioTek Synergy H1 microplate reader according to the manufacturer’s protocol.

#### Western blot analysis

Cells were lysed in RIPA buffer (Thermo, Cat#89900) supplemented with Halt protease inhibitor cocktail (Thermo, Cat# 78429) and 1% n-dodecyl β-D-maltoside (DDM) (Thermo, Cat# BN2005) for solubilization of GPR133 from the plasma membrane. After 15 minutes on ice, lysates were gently sonicated in a water-bath Bioruptor (Diagenode, Cat# UCD-300) at medium power level for 8 cycles of 15 seconds “ON”, 60 seconds “OFF” at 4°C to shear the chromatin. Whole cell lysates were precleared by centrifugation at 15,000 x g for 10 minutes at 4°C. Protein concentrations were determined using the DC protein assay kit II (BioRad, Cat# 5000112). Protein lysates were reduced in Laemmli buffer (BioRad, Cat# 1610747) containing *β*-mercaptoethanol at 37°C for 30 minutes but were not boiled in order to prevent aggregation of the GPR133 transmembrane region. Equal amounts of protein were separated by SDS-PAGE and transferred to 0.2 μm nitrocellulose membranes (BioRad, Cat# 1620112). After blocking the membranes in 2% BSA in TBS-Tween for 1 hour at room temperature, they were simultaneously incubated with multiple primary antibodies of different species (listed in the Key Resources table and **Table S3**) at 4°C overnight and visualized with up to 3 simultaneous fluorescent Alexa Fluor Plus-conjugated secondary antibodies. Images were acquired using the iBrightFL1000 system (Invitrogen). Densitometric quantification of band intensities was conducted in ImageJ.

#### Subcellular fractionation

Adherent cells were gently scraped off from culture dishes in ice-cold PBS in the absence of digestion enzymes or detergents. Subcellular fractionation was conducted using the Plasma Membrane Protein Extraction kit (Abcam, Cat# ab65400) according to the manufacturer’s protocol with slight variations. All steps are conducted at 4°C or on ice. In brief, cells were resuspended in homogenization buffer (Abcam, Cat# ab65400) containing protease inhibitors and were gently broken up using a dounce-homogenizer until >90% of cells were ruptured. Homogenates were centrifuged at 700 x g for 10 minutes and supernatant was transferred to a new tube. The pellet containing nuclei and intracellular organelles of the secretory pathway was kept as “Nuc/ER/Golgi” fraction for this study. The supernatant was centrifuged at 10,000 x g for 30 minutes. The resulting supernatant was kept as “Cytosol/ER” fraction. The resulting pellet was further purified using the two-phase separation of the Plasma Membrane Protein Extraction kit (Abcam, Cat# ab65400) without alterations to the protocol, resulting in the highly pure “plasma membrane” fraction. All fractions were resuspended in 1% n-dodecyl β-D-maltoside (DDM) (Thermo, Cat# BN2005) for better solubilization of the GPR133 transmembrane region. The genomic DNA of the Nuc/ER/Golgi fraction was sheared using a Bioruptor water bath sonicator (Diagenode, Cat# UCD-300) at medium power level for 8 cycles of 15 seconds “ON”, 60 seconds “OFF” at 4°C, and all fractions were precleared by centrifugation at 15,000 x g for 10 minutes. The different subcellular fractions were then either analyzed by Western blot, used as input for affinity purification, or treated with deglycosylating enzyme mixes.

To determine the subcellular distribution of GPR133 fragments (CTF, NTF, H543R Full-length), Western blots of subcellular fractions from GPR133-overexpressing cells were stained using the antibodies targeting the GPR133 C-terminus or N-terminus and analyzed by densitometry in ImageJ. Subcellular distribution was calculated as the protein amount of a fragment in one subcellular fraction, divided by the sum of this fragment’s protein amount across all subcellular fractions (percent of total).

#### Deglycosylation

Whole cell lysates, subcellular fractions, or eluted proteins were treated with endoglycosidase H (EndoHf, NEB, Cat# P0703) or a complete Protein Deglycosylation Mix II (NEB, Cat# P6044) under denaturing conditions for 16 hours at 37°C as recommended by the manufacturer’s protocol. Heating steps beyond 37°C were omitted from the protocol to prevent aggregation of the GPR133 transmembrane region.

#### Affinity purification

Strep-tagII or TwinStrep-tagged GPR133 protein was affinity purified from whole cell lysates, subcellular fractions, or cell culture supernatants using StrepTactin beads (IBA, Cat# 2-1201-002) according to the manufacturer’s protocols. All steps were performed on ice or at 4°C. In brief, input proteins in solution containing 1% n-dodecyl β-D-maltoside (DDM) (Thermo, Cat# BN2005) and protease inhibitor cocktail (Thermo, Cat# 78429) were pretreated with 1/10 volume of 10X Buffer W (IBA, Cat# 2-1003-100) and BioLock (IBA, Cat# 2-0205-250) for 15 minutes on ice to block free biotin. Solutions were then precleared again by centrifugation at 15,000 x g for 15 minutes. Proteins were then incubated with StrepTactin Sepharose beads (IBA, Cat# 2-1201-002) rotating at 4°C overnight. The following day, StrepTactin beads were pelleted and washed 5 times in a large excess of 1X Buffer W containing 0.1% DDM and protease inhibitors rotating at 4°C for 30 minutes (e.g. 20 μL beads washed with 1 mL buffer). Proteins were eluted from StrepTactin beads in 3 consecutive elutions with 1X Buffer E (IBA, Cat# 2-1000-025) containing 1% DDM and supplemented with desthiobiotin (IBA, Cat# 2-1000-001) to a total concentration of 10 mM. Elutions were pooled and analyzed by Western blot or subjected to deglycosylation as described above.

#### Immunofluorescent staining and microscopy

Cells cultured on poly-L-ornithine (Sigma, Cat# P4957) and laminin (Thermo Fisher, Cat# 23017015) pretreated slides were washed gently with ice cold PBS and fixed with 4% paraformaldehyde (PFA, Sigma-Aldrich, Cat# P6148) for 30 minutes at room temperature. For permeabilizing conditions, all following steps were conducted in the presence of 0.1% Triton X-100; for non-permeabilizing steps, no detergent was added until the nuclear counter staining with DAPI. Cells were blocked with 10% bovine serum albumin (BSA) in PBS for 1 hour at room temperature and stained with primary antibody mixes in 1% BSA in PBS at 4°C overnight. Antibodies and the concentrations used are detailed in **Table S3**. Primary antibody staining was visualized by staining with Alexa Fluor Plus conjugated secondary antibodies at a concentration of 1:1,000 for 1 hour at room temperature. Nuclei were counterstained with 500 ng/mL 4′,6-diamidino-2-phenylindole (DAPI) for 10 minutes at room temperature. Confocal laser scanning microscopy was conducted on a Zeiss LSM700 and images were analyzed and exported from ImageJ software.

#### Protein decay and half-life analysis

HEK293T cells were transfected with various GPR133 overexpression constructs using Lipofectamine 2000 (Invitrogen, Cat# 11668-019) and each split into multiple 6-well plates after 24 hours. Forty-eight hours after transfection all wells were treated with 280 μg/mL cycloheximide (Sigma-Aldrich, Cat# 239765). The “0 hour” control condition was harvested right away, other conditions were harvested 2, 4, 8, 12, and 24 hours later. Cells were washed once with PBS and whole cell lysates were prepared and analyzed by Western blot as described above. For each resulting Western blot lane, the amounts of GPR133 fragments as detected by the antibodies targeting the C-terminus or N-terminus were analyzed by densitometry on ImageJ. Protein amounts of the GPR133 fragments normalized to their “0-hours” control conditions were plotted as a function of time in GraphPad Prism (modeled one phase decay curves) to determine the protein half-life.

#### Overexpression plasmids and molecular cloning

All overexpression plasmids are based on the lentiviral vector pLVX-EF1alpha-mCherry-N1 (Takara, Cat# 631986). Codon-optimized cDNA encoding for wild-type human GPR133 (ADGRD1) was obtained from Axxam (Milan, Italy) and subcloned into the pLVX vector using Gibson Assembly, replacing the puromycin resistance (PuroR) gene. Human PAR1 receptor cDNA was obtained from Sino Biologicals (Cat#, HG13535-NM). hPAR1-GPR133 fusion constructs as well as all other modifications to the GPR133 overexpression construct were subcloned by Gibson Assembly.

#### Structural prediction

The 3D protein structure of GPR133 was predicted and modeled using the Protein Homology/analogY Recognition Engine V 2.0 (Phyre2) web portal (http://www.sbg.bio.ic.ac.uk/~phyre2/html/page.cgi?id=index) (Kelley et al., 2015). The analysis was run in the “Intensive” mode using the following amino acid sequence as input: MEKLLRLCCWYSWLLLFYYNFQVRGVYSRSQDHPGFQVLASASHYWPLENVDGIHELQDTTGDIVEGKVNKGIYLKEEKGVTLLYYGRYNSSCISKPEQCGPEGVTFSFFWKTQGEQSRPIPSAYG GQVISNGFKVCSSGGRGSVELYTRDNSMTWEASFSPPGPYWTHVLFTWKSKEGLKVYVNGTLSTSDPSGKVSRDYGESNVNLVIGSEQDQAKCYENGAFDEFIIWERALTPDEIAMYFTAAIGKHALLSSTLPSLFMTSTASPVMPTDAYHPIITNLTEERKTFQSPGVILSYLQNVSLSLPSKSLSEQTALNLTKTFLKAVGEILLLPGWIALSEDSAVVLSLIDTIDTVMGHVSSNLHGSTPQVTVEGSSAMAEFSVAKILPKTVNSSHYRFPAHGQSFIQIPHEAFHRHAWSTVVGLLYHSMHYYLNNIWPAHTKIAEAMHHQDCLLFATSHLISLEVSPPPTLSQNLSGSPLITVHLKHRLTRKQHSEATNSSNRVFVYCAFLDFSSGEGVWSNHGCALTRGNLTYSVCRCTHLTNFAILMQVVPLELARGHQVALSSISYVGCSLSVLCLVATLVTFAVLSSVSTIRNQRYHIHANLSFAVLVAQVLLLISFRLEPGTTPCQVMAVLLHYFFLSAFAWMLVEGLHLYSMVIKVFGSEDSKHRYYYGMGWGFPLLICIISLSFAMDSYGTSNNCWLSLASGAIWAFVAPALFVIVVNIGILIAVTRVISQISADNYKIHGDPSAFKLTAKAVAVLLPILGTSWVFGVLAVNGCAVVFQYMFATLNSLQGLFIFLFHCLLNSEVRAAFKHKTKVWSLTSSSARTSNAKPFHSDLMNGTRPGMASTKLSPWDKSSHSAHRVDLSAV. Phyre2 reports 93% of the residues to be modeled with >90% confidence. The resulting pdb file was visualized in PyMOL in a cartoon ribbon format, domains were pseudocolored, and the *Stachel* region was highlighted as a surface model. This predicted structure has not been experimentally validated and only serves as an approximate visual guide.

## QUANTIFICATION AND STATISTICAL ANALYSIS

All statistical comparisons were conducted in GraphPad Prism (v9). Population statistics were represented as mean ± SEM throughout. Statistical significance was calculated using Students t-test; and one-way or two-way analysis of variance (ANOVA) with *post hoc* Tukey’s multiple comparisons test. The threshold of significance was set at P<0.05.

## SUPPLEMENTAL FIGURE CAPTIONS

**Figure S1.**
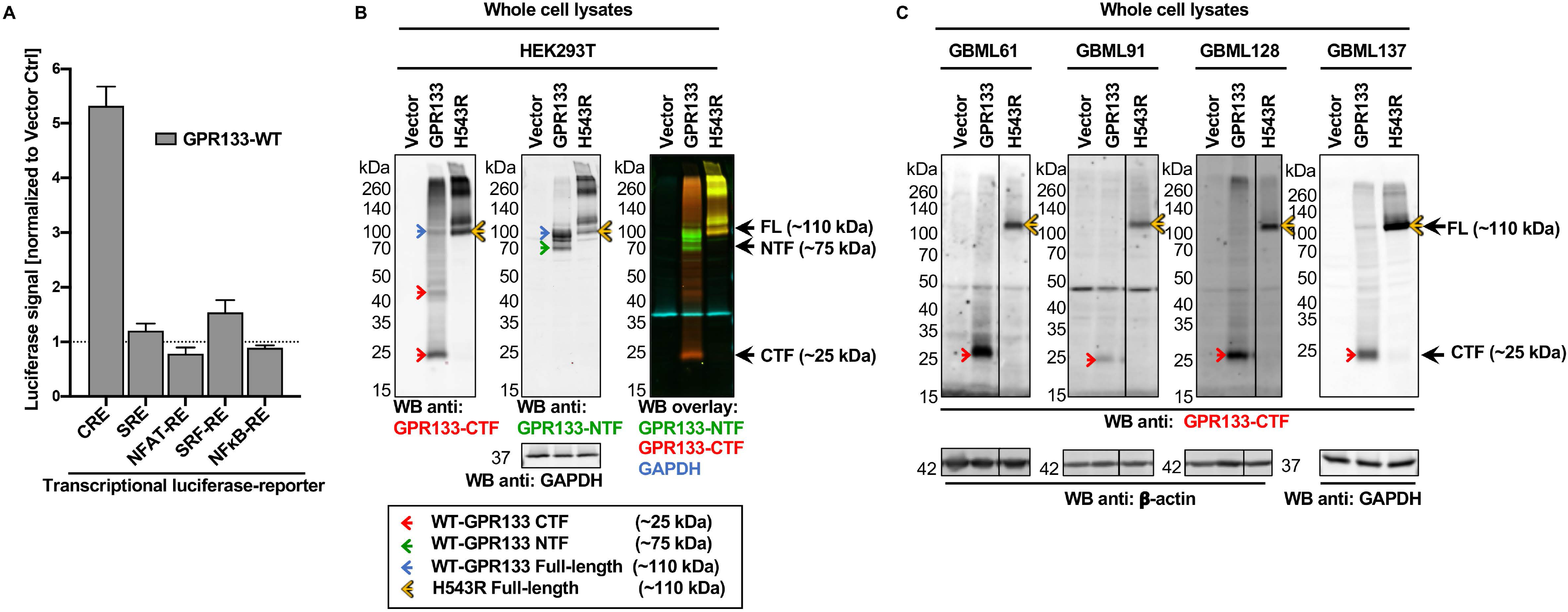
Wild-type GPR133 is cleaved in patient-derived GBM and HEK293T cells and signals through Gα_S_ -mediated cAMP increase. **A)** HEK293T cells were co-transfected with WT GPR133 and luciferase reporters for various known G protein-mediated signaling pathways. Luciferase signals are expressed as mean ± SEM of normalized fold change over vector control. GPR133 overexpression in combination with a CRE-Luciferase reporter plasmid resulted in a robust increase in luciferase activity, confirming cAMP-mediated signaling as its main canonical signaling pathway (Luciferase fold change over vector: CRE: 5.32 ± 0.35; SRE: 1.21 ± 0.13; NFAT: 0.79 ± 0.11; SRF-RE: 1.54 ± 0.22; NFκB: 0.89± 0.04; ANOVA: F_(4, 25)_ = 76.45; P<0.0001; n=5-8 per reporter). **B)** Western blot analysis of whole cell lysates from HEK239T cells overexpressing WT or H543R mutant GPR133 (used as inputs to generate Figure 1E). HEK293T cells were transfected with an empty vector control, WT, or H543R mutant GPR133, lysed, and analyzed by Western blot. Membranes were co-stained with an antibody detecting the GPR133 CTF (left panel, red staining in WB overlay) and an antibody against the GPR133 NTF (middle panel, and green staining in WB overlay). The C-terminal antibody detects the cleaved WT GPR133 CTF monomer at ∼25 kDa and putative dimer at ∼48 kDa (red arrows), as well as an uncleaved WT GPR133 at ∼110 kDa (blue arrows). The N-terminal antibody detects the cleaved WT NTF at ∼75 kDa (green arrow) as well as the uncleaved WT GPR133 at ∼110 kDa (blue arrows). Both the CTF- and NTF-targeting antibodies detect the uncleaved H543R mutant full-length receptor at ∼110 kDa (yellow arrows). **C)** Four separate patient-derived GBM cultures were transduced via lentivirus (GBML61, 91, 128) or by transfection (GBML137) with either an empty vector control, WT, or H543R mutant GPR133. Whole cell lysates were analyzed by Western blot using an antibody targeting the GPR133 CTF. In all four cultures, WT GPR133 is detected almost entirely cleaved as CTF band at ∼25 kDa (red arrows), while H543R mutant GPR133 is detected as uncleaved band at ∼110 kDa (yellow arrows). GBML128 is the corresponding input sample used to generate Figure 1F. Membranes were counterstained against β-actin or GAPDH as loading control (lower panels).

**Figure S2.**
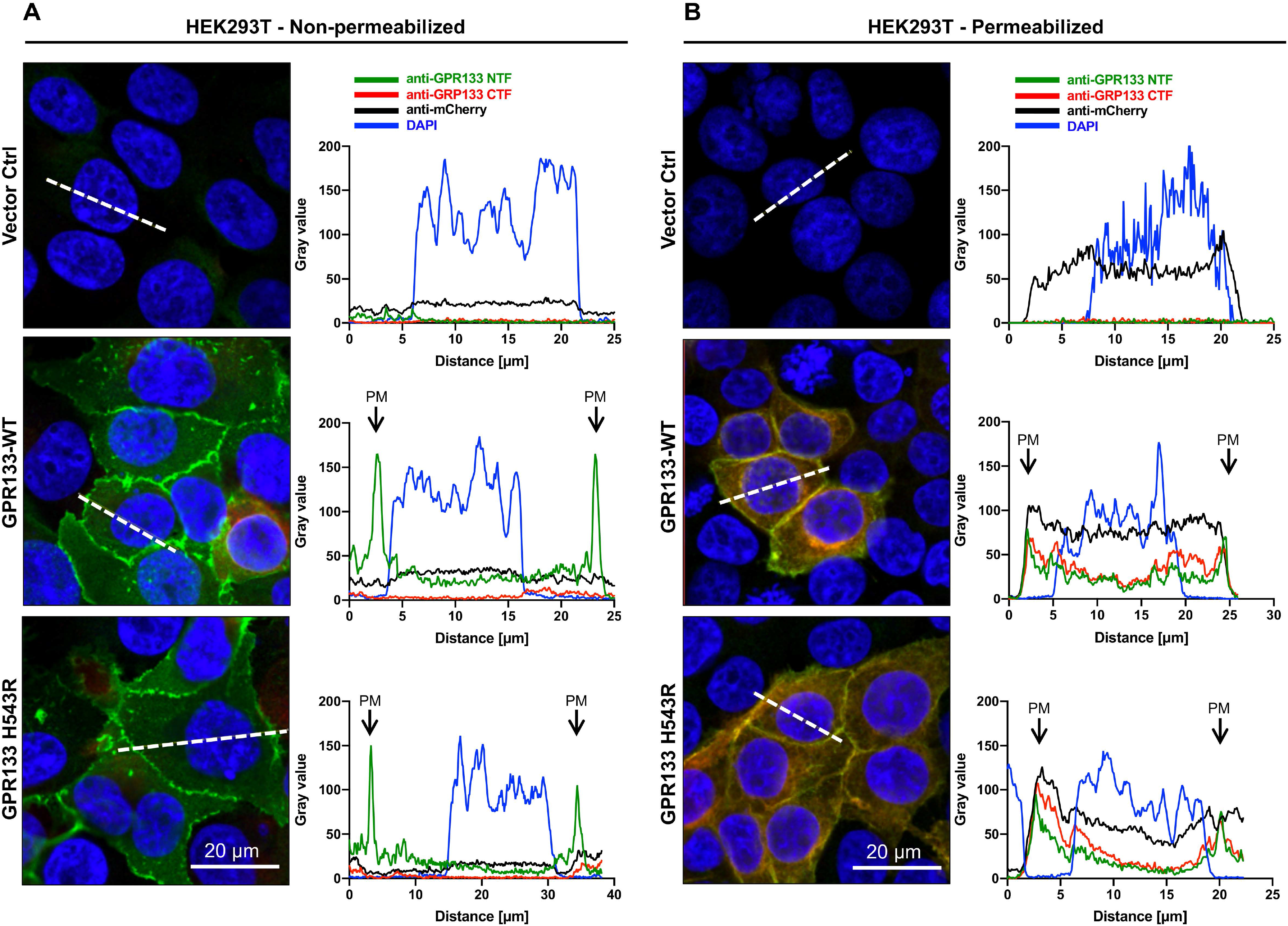
WT and H543R mutant GPR133 localize to the plasma membrane. **A, B**) Representative confocal microscopy micrographs of HEK293T cells overexpressing GPR133 demonstrate that intramolecular cleavage is not required for plasma membrane localization. Cells were transfected with either a vector control, WT, or H543R mutated GPR133, fixed and fluorescently stained under non-permeabilizing (**A**) or permeabilizing (**B**) conditions against the GPR133 CTF, GPR133 NTF, and transfection marker mCherry (not included in the composite image), as described for Figure 2. The fluorescence intensity across all channels was measured along a virtual 2D cross section through the cells (shown by the white dotted line) and plotted as a function of distance in micrometers. Transfected cells demonstrate distinct intensity peaks for GPR133 staining at the plasma membrane for both the WT and the uncleaved H543R mutant GPR133. Nuclei were counterstained with DAPI. Scale bars, 20 µm.

**Figure S3.**
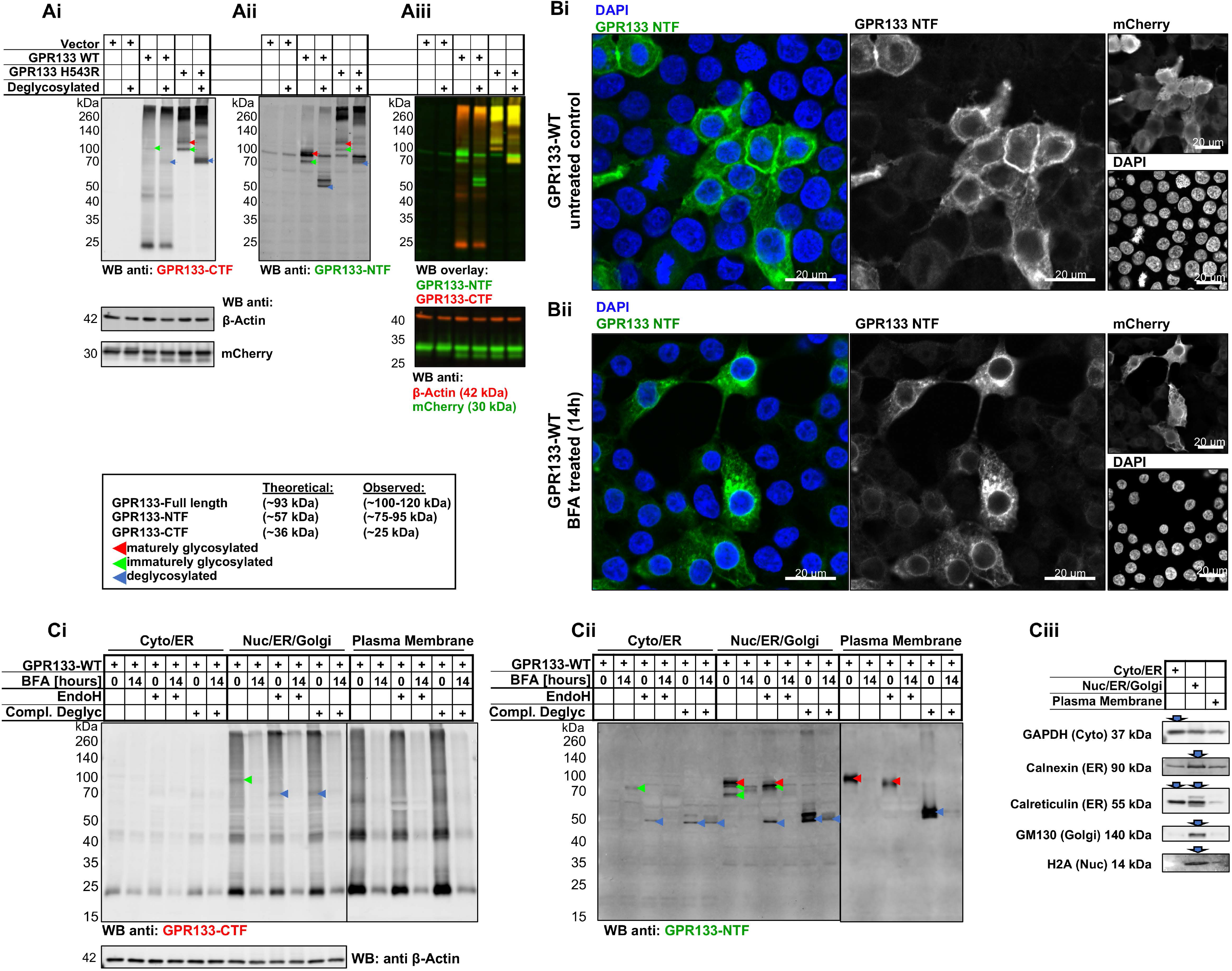
Intramolecular cleavage of GPR133 occurs in the ER and prior to mature glycosylation. Presumed maturely glycosylated, immaturely glycosylated, and completely deglycosylated forms of GPR133 are marked with red, green, and blue arrowheads respectively, throughout. **A)** Whole cell lysates of HEK293T cells overexpressing a vector control, WT, or H543R mutated GPR133 were subjected to complete deglycosylation or control treatment and analyzed by Western blot. Membrane was counterstained against β-actin and mCherry as loading controls (lower panels). **B)** Representative micrographs of HEK293T cells overexpressing WT GPR133 and treated with DMSO control (**Bi**) or Brefeldin A (BFA) for 14 hours (**Bii**). Cells were fixed, permeabilized, fluorescently stained against the GPR133 NTF, and analyzed by confocal laser scanning microscopy. Untreated control cells demonstrate plasma membrane localization of GPR133, while BFA treated cells demonstrate the brightest GPR133 staining in the perinuclear region. mCherry is co-expressed on all vectors used in this study and is included in the single-channel panels as transfection control but is not included in the composite panels. Nuclei were counterstained with DAPI. Scale bars denote 20 µm. **C)** HEK293T cells overexpressing GPR133 were treated with either a DMSO control or 3 μg/mL brefeldin A (BFA) for 14 hours to block ER-to-Golgi transport. Cropped untreated lanes are depicted separately in Figure 3D. Cell lysates were subjected to subcellular fractionation followed by treatment with either endoglycosidase H (EndoH) or a complete enzymatic deglycosylation mix. Note that the uncleaved form of WT GPR133 is sensitive to EndoH treatment (**Ci**), and that the cleaved NTF exists as both immaturely glycosylated EndoH-sensitive, and maturely glycosylated EndoH-insensitive forms (**Cii**). Complete deglycosylation confirms that all observed isoforms of the NTF are the same protein with varying degrees of glycosylation (blue arrowheads). **Ciii**) Subcellular compartment markers validate enrichment of the respective subcellular fractions (same membranes also depicted in Figure 3Diii).

**Figure S4.**
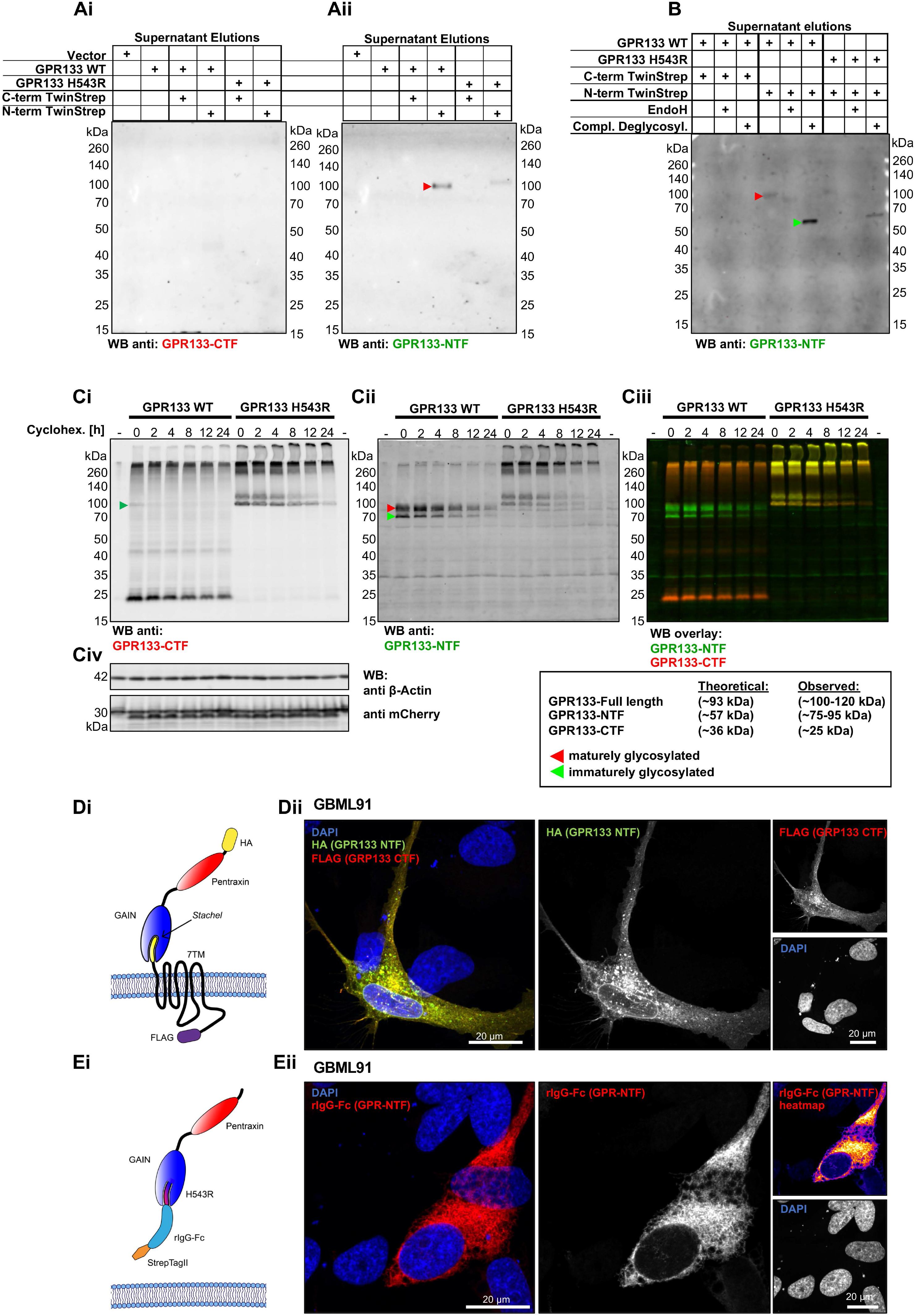
The GPR133 NTF dissociates from the CTF at the plasma membrane. **A)** Soluble GPR133 NTF is detected in precleared cell culture supernatants. Supernatants from HEK293T cells overexpressing various tagged GPR133 constructs were harvested, precleared, and used as inputs for affinity purification. Western blot membranes depict the resulting elutions stained against the GPR133 CTF (**Ai**) and NTF (**Aii**) respectively. Multiple independent repeats are depicted and quantified in main Figure 4E. **B)** Deglycosylation of the supernatant elutions depicted in Figure 4Eii confirms the detected band to be the cleaved GPR133 NTF by molecular weight (∼57 kDa after complete deglycosylation). **C)** Representative Western blot membranes showing the time-course of GPR133 fragments after blocking protein synthesis with cycloheximide. HEK293T cells overexpressing WT or H543R mutant GPR133 were treated with 280 μg/mL cycloheximide and lysed after varying amounts of time. Resulting whole cell lysates were analyzed by Western blot and GPR133 fragments were quantified by densitometry. The highly stable proteins β-Actin and mCherry were used as loading control (**Civ**). **D)** GPR133 is not detected on GBM cells adjacent to ectopic GPR133 overexpressing cells. Patient-derived GBM cultures (GBML91) were sparsely transfected with an ectopic overexpression construct of C- and N-terminally tagged GPR133 (schematic **Di**). Cells were fixed, permeabilized, and stained against the C-terminal FLAG-tag and N-terminal HA-tag of the ectopic GPR133 and analyzed by confocal microscopy. A representative micrograph is depicted in **Dii**. While the tagged ectopic GPR133 NTF was detected on the infected GBM cells (marked by staining for the GPR133-CTF), no additional staining was observed on adjacent cells. Nuclei were counterstained with DAPI. Scale bars, 20 µm. **E)** A tagged, secreted form of the GPR133 NTF containing a H543R mutation and the *Stachel* region for structural integrity was overexpressed and analyzed by confocal microscopy as detailed for panel **D**. No ectopic secreted GPR133-NTF is detected on adjacent cells. Heatmap depiction of single channel staining of the tagged NTF is included for higher visual sensitivity. Nuclei were counterstained with DAPI. Scale bars, 20 µm.

**Figure S5.**
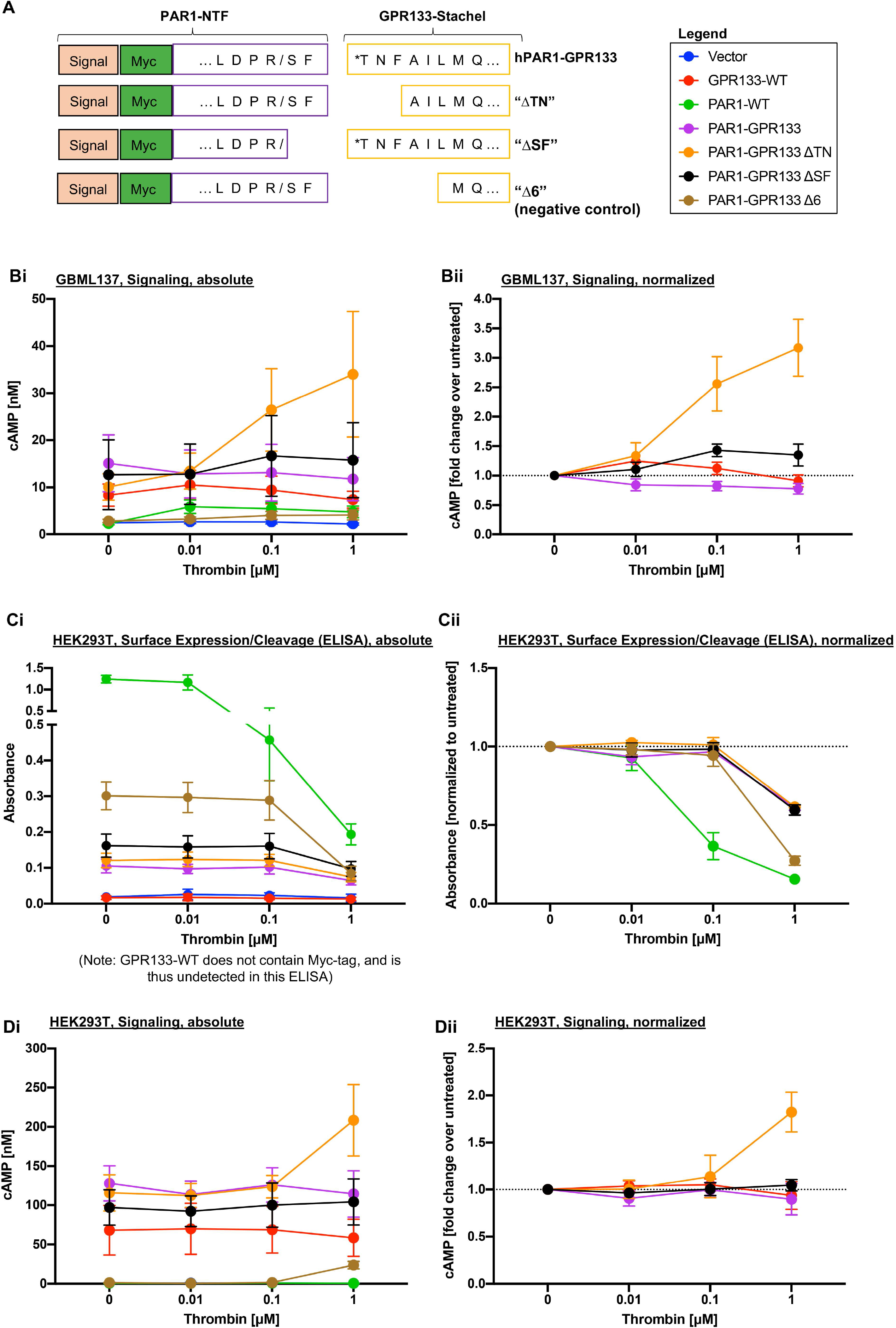
NTF shedding at the plasma membrane increases canonical signaling of a hybrid hPAR1-GPR133 receptor. **A)** Schematic of the hPAR1-GPR133 fusion construct design. The first two amino acids subsequent to the thrombin recognition site (SF) were reported to be critical for thrombin-mediated cleavage of the human PAR1-NTF. Mathiasen and colleagues proposed to replace the first three amino acids of the *Stachel* sequence (TNF) with these residues in their study on ADGRL1 to create a fusion construct both capable of cleavage and canonical receptor signaling (“ΔTN”) (Mathiasen et al., 2020). Multiple variations of the fusion between hPAR1-NTF and GPR133-CTF were included in our study. The “Δ6” construct is lacking the first 6 residues of the *Stachel* sequence and functions as a negative control. “/” and “*” mark the site of thrombin-mediated cleavage and the endogenous GPR133 cleavage site respectively. **Bi**) Patient-derived GBM cells (GBML137) overexpressing either WT GPR133 or the various PAR1-GPR133 fusion were exposed to varying concentrations of thrombin for 30 minutes in the presence of 1 mM IBMX, and intracellular cAMP levels were assessed by HTRF assays. Data is depicted as mean ± SEM of absolute cAMP levels. Thrombin-mediated dissociation of the NTF significantly increased canonical GPR133 signaling in the hPAR1-GPR133-ΔTN fusion construct, but none of the other constructs (Two-way ANOVA: construct F_(6,56)_=7.83, P<0.0001, thrombin F_(3,56)_=0.98, P=0.41, interaction of factors F_(18,56)_=0.79, P=0.69; Tukey’s multiple comparisons hPAR1-GPR133-ΔTN 0 µM vs 1 µM thrombin P<0.01; 0.01 µM vs 1 µM thrombin P<0.05; all other comparisons within each construct are not significant; n=3 independent experiments with technical triplicates). **Bii**) Data from panel **Bi** normalized to untreated condition expressed as mean ± SEM (Two-way ANOVA: construct F_(3,32)_=27.18, P<0.0001, thrombin F_(3,32)_=7.61, P<0.001, interaction of factors F_(9,32)_=7.32, P<0.0001; Tukey’s multiple comparisons hPAR1-GPR133-ΔTN 0 µM vs 0.1 µM thrombin P<0.0001; 0 µM vs 1 µM thrombin P<0.0001; 0.01 µM vs 0.1 µM thrombin P<0.001; 0.01 µM vs 1 µM thrombin P<0.0001; all other comparisons within each construct are not significant; n=3 independent experiments with technical triplicates). **Ci)** Cell surface ELISA detects the thrombin-mediated dissociation of the Myc-tagged PAR1-NTF in the full-length PAR1 (positive control) and all PAR1-GPR133 fusion protein variants (Two-way ANOVA: construct F_(6,56)_=117.8, P<0.0001, thrombin F_(3,56)_=27.6, P<0.0001, interaction of factors F_(18,56)_=15,7, P<0.0001; n=3 independent experiments with technical triplicates). Cells were exposed to thrombin for 30 minutes, fixed, and stained against the N-terminal Myc-tag under non-permeabilizing conditions. Of note, the WT GPR133 and the vector control do not contain a Myc-tag and are therefore not detected. ELISA absorbance is displayed in non-normalized arbitrary units as mean ± SEM. **Cii**) Data from panel **Ci** normalized to untreated condition expressed as mean ± SEM (Two-way ANOVA: construct F_(4,40)_=38.7, P<0.0001, thrombin F_(3,40)_=206.0, P<0.0001, interaction of factors F_(12,40)_=13.8, P<0.0001; n=3 independent experiments with technical triplicates). **Di)** HEK293T cells overexpressing either WT GPR133 or the various PAR1-GPR133 fusion constructs were exposed to varying concentrations of thrombin for 30 minutes in the presence of 1 mM IBMX, and intracellular cAMP levels were assessed by HTRF assays. Data is depicted as mean ± SEM of absolute cAMP levels. Thrombin-mediated dissociation of the NTF significantly increased canonical GPR133 signaling in the hPAR1-GPR133-ΔTN fusion construct, but none of the other constructs (Two-way ANOVA: construct F_(6,52)_=30.9, P<0.0001, thrombin F_(3,52)_=0.86, P=0.47, interaction of factors F_(18,52)_=0.76, P=0.78; Tukey’s multiple comparisons hPAR1-GPR133-ΔTN 0 µM vs 1 µM thrombin P<0.05; 0.01 µM vs 1 µM thrombin P<0.05; 0.1 µM vs 1 µM thrombin P<0.05; all other comparisons not significant; n=3 independent experiments with technical triplicates). **Dii**) Data from panel **Di** normalized to untreated condition expressed as mean ± SEM (Two-way ANOVA: construct F_(3,32)_=5.87, P<0.003, thrombin F_(3,32)_=2.72, P=0.06, interaction of factors F_(9,32)_=3.75, P<0.003; Tukey’s multiple comparisons hPAR1-GPR133-ΔTN 0 µM vs 1 µM thrombin P<0.0001; 0.01 µM vs 1 µM thrombin P<0.0001; 0.1 µM vs 1 µM thrombin P<0.0006; all other comparisons within each construct are not significant; n=3 independent experiments with technical triplicates).

---

**Table S1.** Genetic background of patient-derived GBM cultures

| **Patient-derived culture** | **Gender** | **IDH status** | **MGMT promoter methylation** | **EGFR amplification** | **Oncomine focus assay** |
| --- | --- | --- | --- | --- | --- |
| **GBML61** | F | wild-type | not methylated | amplified | FGFR3 F386L mutation, JAK3 V722I mutation |
| **GBML91** | F | wild-type | methylated | not amplified | no mutations |
| **GBML128** | F | wild-type | methylated | not tested | no mutations |
| **GBML137** | F | wild-type | not methylated | not tested | *PDGFRA* amplification, *CDK4* amplification |

IDH, isocitrate dehydrogenase; MGMT, O<sup>6</sup> methylguanine DNA methyltransferase; EGFR, epidermal growth factor receptor.

**Table S2.** Genes sequenced in NYU Oncomine focus assay

|  |  |
| --- | --- |
| **Mutations** | AKT1, ALK, AR, BRAF, CDK4, CTNNB1, DDR2, EGFR, ERBB2, ERBB3, ERBB4, ESR1, FGFR2, FGFR3, GNA11, GNAQ, HRAS, IDH1, IDH2, JAK1, JAK2, JAK3, KIT, KRAS, MET, MTOR, NRAS, PDGFRA, PIK3CA, RAF1, RET, ROS1, SMO |
| **CNV** | ALK, AR, BRAF, CCDN1, CDK4, CDK6, EGFR, ERBB2, FGFR1, FGFR2, FGFR3, FGFR4, KIT, KRAS, MET, MYC, MYCN, PDGFRA, PIK3CA |
| **Fusion drivers** | ABL1, AKT3, ALK, AXL, BRAF, ERG, ETV1, ETV4, ETV5, EGFR, ERBB2, FGFR1, FGFR2, FGFR3, MET, NTRK1, NTRK2, NTRK3, PDGFRA, PPARG, RAF1, RET, ROS1 |

CNV, copy number variation.

**Table S3.** Antibodies used in this study

| **Antibody target** | **Host Species** | **Dilution WB** | **Dilution IF** | **Manufacturer** | **Cat. No.** |
| --- | --- | --- | --- | --- | --- |
| GPR133-CTF | Rabbit | 1:1,000 | 1:200 | Sigma-Aldrich | HPA042395 |
| GPR133-NTF | Mouse monoclonal | 1:20,000 | 1:8,000 | Not commercially available (Bayin et al. 2016; Frenster et al., 2020) | Clone "8E3E8" |
| Beta-Actin (AC15) | Mouse monoclonal | 1:1000 | - | Thermo Fisher | AM4302 |
| GAPDH | Goat | 1:1,000 | - | Thermo Fisher | PA1-9046 |
| mCherry | Chicken | 1:2,000 | 1:1,000 | Abcam | ab205402 |
| Calnexin | Rabbit | 1:1,000 | - | Cell Signaling | 2433S |
| Calreticulin (EPR3924) | Recombinant Rabbit | 1:2,000 | - | Abcam | ab92516 |
| GM130 (EP892Y) | Recombinant Rabbit | 1:2,000 | - | Abcam | ab52649 |
| Pan Cadherin (EPR1792Y) | Recombinant Rabbit | 1:10,000 | - | Abcam | ab51034 |
| Histone H2A (L88A6) | Mouse monoclonal | 1:2,000 | - | Cell Signaling | 3636S |
| ATP1A3 (XVIF9-G10) | Mouse monoclonal | 1:1,000 | - | Thermo Fisher | MA3-915 |
| Myc-tag (9B11) | Mouse monoclonal | 1:1000 | 1:500 | Cell Signaling | 2276S |
| FLAG-tag (M2) | Rabbit | 1:1000 | 1:4,000 | Sigma-Aldrich | F7425 |
| HA-tag | Mouse monoclonal | 1:1000 | 1:2,000 | Sigma-Aldrich | H3663 |
| anti-Mouse IgG Alexa Fluor Plus 488 | Donkey | 1:5,000 | 1:1,000 | Thermo Fisher | A32766 |
| anti-Rabbit IgG Alexa Fluor Plus 555 | Donkey | 1:5,000 | 1:1,000 | Thermo Fisher | A32794 |
| anti-Goat IgG Alexa Fluor 647 | Donkey | 1:5,000 | 1:1,000 | Thermo Fisher | A21447 |
| anti-Rabbit<br>Alexa Fluor<br>647 | Donkey | 1:5,000 | 1:1,000 | Thermo Fisher | CA31573 |
| anti-Chicken<br>IgY Alexa Fluor<br>Plus 555 | Goat | 1:5,000 | 1:1,000 | Thermo Fisher | A32932 |

## KEY RESOURCES TABLE

| REAGENT or RESOURCE | SOURCE | IDENTIFIER |
| --- | --- | --- |
| **Antibodies** |  |  |
| Rabbit polyclonal anti-GPR133-CTF | Sigma-Aldrich | Cat# HPA042395 |
| Mouse monoclonal anti-GPR133-NTF | Not commercially available (Bayin et al. 2016) (Frenster et al., 2020) | Clone "8E3E8" |
| Mouse monoclonal anti-Beta-Actin (AC15) | Thermo Fisher | Cat# AM4302 |
| Goat polyclonal anti-GAPDH | Thermo Fisher | Cat# PA1-9046 |
| Chicken polyclonal anti-mCherry | Abcam | Cat# ab205402 |
| Rabbit polyclonal anti-Calnexin | Cell Signaling | Cat# 2433S |
| Recombinant anti-Calreticulin (EPR3924) | Abcam | Cat# ab92516 |
| Recombinant anti-GM130 (EP892Y) | Abcam | Cat# ab52649 |
| Recombinant anti-Pan Cadherin (EPR1792Y) | Abcam | Cat# ab51034 |
| Mouse monoclonal anti-Histone H2A (L88A6) | Cell Signaling | Cat# 3636S |
| Mouse monoclonal anti-ATP1A3 (XVIF9-G10) | Thermo Fisher | Cat# MA3-915 |
| Mouse monoclonal anti-Myc-tag (9B11) | Cell Signaling | Cat# 2276S |
| Rabbit polyclonal anti-FLAG-tag (M2) | Sigma-Aldrich | Cat# F7425 |
| Mouse monoclonal anti-HA-tag | Sigma-Aldrich | Cat# H3663 |
| Secondary Donkey anti-Mouse Alexa Fluor Plus 488 | Thermo Fisher | Cat# A32766 |
| Secondary Donkey anti-Rabbit Alexa Fluor Plus 555 | Thermo Fisher | Cat# A32794 |
| Secondary Donkey anti-Goat Alexa Fluor 647 | Thermo Fisher | Cat# A21447 |
| Secondary Donkey anti-Rabbit Alexa Fluor 647 | Thermo Fisher | Cat# A31573 |
| Secondary Goat anti-Chicken IgY Alexa Fluor Plus 555 | Thermo Fisher | Cat# A32932 |
| Secondary Chicken anti-Mouse HRP-conjugated | Thermo Fisher | Cat# A15975 |
| **Bacterial and Virus Strains** |  |  |
| **Biological Samples** |  |  |
| **Chemicals, Peptides, and Recombinant Proteins** |  |  |
| Recombinant human Thrombin | Sigma-Aldrich | T9326-150UN |
| Protein Deglycosylation Mix II | NEB | P6044 |
| Endoglycosidase Hf | NEB | P0703 |
| Cycloheximide - CAS 66-81-9 | Sigma-Aldrich | 239765-1ML |
| Brefeldin A Solution (1000X) | Invitrogen | 00-4506-51 |
| 3-Isobutyl-1-methylxanthine (IBMX) | Sigma-Aldrich | I7018-100MG |
| DDM (n-dodecyl β-D-maltoside) (10%) | Thermo Scientific | BN2005 |
| Neurobasal Medium | Gibco | 21103049 |
| EGF | R&D | 236-EG-01M |
| bFGF | R&D | 233-FB-01M |
| GlutaMAX Supplement | Gibco | 35050061 |
| MEM Non-Essential Amino Acids | Gibco | 11140050 |
| N2-Supplement | Gibco | 17-502-049 |
| B27-Supplement | Gibco | 12587010 |
| Poly-L-Ornithine solution | Sigma-Aldrich | P4957-50ml |
| Laminin for Cell Culture | Fisher | 23017015 |
| Lipofectamine 2000 | Invitrogen | 11668-019 |
| Lipofectamine Stem Reagent (used for GBM cells) | Thermo Fisher | STEM00008 |
| Accutase | Innovative Cell Technologies | AT104 |
| StrepTactin Sepharose resin | IBA | 2-1201-002 |
| Gravity flow Strep-Tactin Sepharose column | IBA | 2-1202-001 |
| 10x Buffer E; Strep-Tactin Elution Buffer with Desthiobiotin | IBA | 2-1000-025 |
| 10x Buffer R; Strep-Tactin Regeneration Buffer with HABA | IBA | 2-1002-100 |
| 10x Buffer W; Strep-Tactin Wash Buffer | IBA | 2-1003-100 |
| BioLock Biotin blocking solution | IBA | 2-0205-250 |
| Desthiobiotin | IBA | 2-1000-001 |
| Critical Commercial Assays |  |  |
| cAMP Gs Dynamic kit (HTRF) | CisBio | 62AM4PEC |
| Plasma membrane extraction kit | Abcam | ab65400 |
| Bright-Glo Luciferase Assay System | Promega | E2650 |
| ELISA: TMB Stabilized Chromogen | Thermo Fisher | SB02 |
| ELISA: Stop Solution | Thermo Fisher | SS04 |
| Deposited Data |  |  |
| Experimental Models: Cell Lines |  |  |
| Lenti-X 293T Cell Line | Takara | 632180 |
| Patient-derived GBM cultures | Derived in our lab, not commercially available | GBML61/91/128/137 |
| Experimental Models: Organisms/Strains |  |  |
| Oligonucleotides |  |  |
| Recombinant DNA |  |  |
| CRE-Luciferase (pGL4.29[luc2P/CRE/Hygro]) | Promega | E8471 |
| SRE-Luciferase (pGL4.33[luc2P/SRE/Hygro]) | Promega | E1340 |
| NFAT-RE-Luciferase (pGL4.30[luc2P/NFAT-RE/Hygro]) | Promega | E8481 |
| SRF-Luciferase (pGL4.34[luc2P/SRF/Hygro]) | Promega | E1350 |
| NFkB-RE-Luciferase (pGL4.32[luc2P/NFkB-RE/Hygro]) | Promega | E8491 |
| Thrombin Receptor cDNA ORF Clone, Human, N-Myc tag | SinoBiologicals | HG13535-NM |
| pLVX-EF1alpha-mCherry-N1 Vector | Takara | 631986 |
| pLVX_EF1a-mCherry_PGK-GPR133-WT | This paper | N/A |
| pLVX_EF1a-mCherry_PGK-GPR133-H543R | This paper | N/A |
| pLVX_EF1a-mCherry_PGK-GPR133-WT-N-terminal_TwinStrep | This paper | N/A |
| pLVX_EF1a-mCherry_PGK-GPR133-WT-C-terminal_TwinStrep | This paper | N/A |
| pLVX_EF1a-mCherry_PGK-GPR133-H543R-N-terminal_TwinStrep | This paper | N/A |
| pLVX_EF1a-mCherry_PGK-GPR133-H543R-C-terminal_TwinStrep | This paper | N/A |
| pLVX_EF1a-mCherry_PGK-HA-GPR133-FLAG | This paper | N/A |
| pLVX_EF1a-mCherry_PGK-GPR133-NTF-H543R-rlgG-Fc-StrepTagII | This paper | N/A |
| pLVX_EF1a-mCherry_PGK-hPAR1-CTF-GPR133-NTF | This paper | N/A |
| pLVX_EF1a-mCherry_PGK-hPAR1CTF-dTNF-GPR133-NTF | This paper | N/A |
| pLVX_EF1a-mCherry_PGK-hPAR1CTF-dSF-GPR133-NTF | This paper | N/A |
| pLVX_EF1a-mCherry_PGK-hPAR1CTF-d5-GPR133-NTF | This paper | N/A |
| Software and Algorithms |  |  |
| ImageJ | Schneider et al., 2012 | https://imagej.nih.gov/ij/ |
| Phyre2 | Kelley LA et al., 2015 | http://www.sbg.bio.ic.ac.uk/phyre2/html/page.cgi?id=index |
| PyMOL |  | https://pymol.org/2/ |
| Other |  |  |

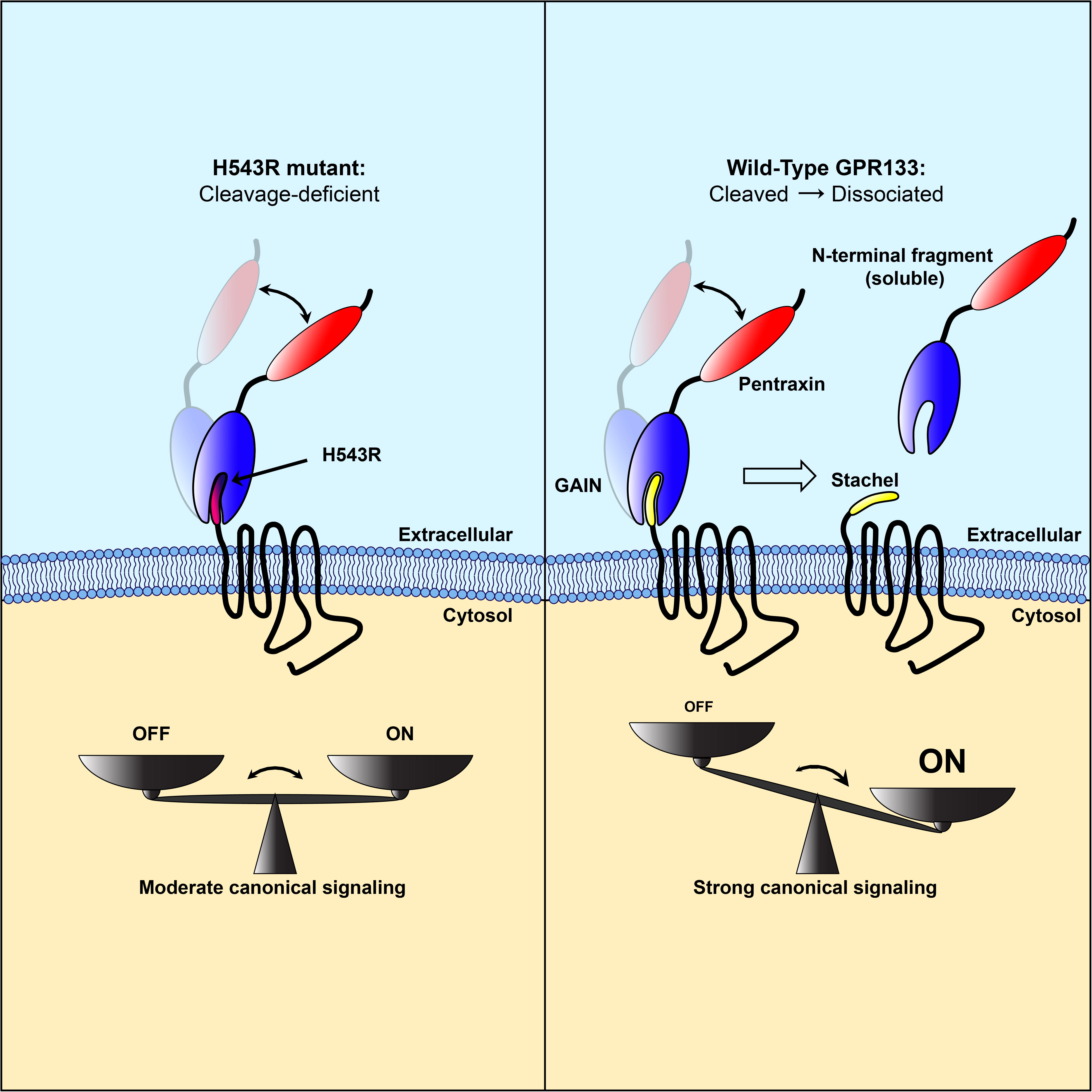

## REFERENCES

Abe, J., Fukuzawa, T., and Hirose, S. (2002). Cleavage of Ig-Hepta at a “SEA” module and at a conserved G protein-coupled receptor proteolytic site. J Biol Chem 277, 23391–23398.

Arac, D., Boucard, A.A., Bolliger, M.F., Nguyen, J., Soltis, S.M., Sudhof, T.C., and Brunger, A.T. (2012). A novel evolutionarily conserved domain of cell-adhesion GPCRs mediates autoproteolysis. EMBO J 31, 1364–1378.

Bayin, N.S., Frenster, J.D., Kane, J.R., Rubenstein, J., Modrek, A.S., Baitalmal, R., Dolgalev, I., Rudzenski, K., Scarabottolo, L., Crespi, D., et al. (2016). GPR133 (ADGRD1), an adhesion G-protein-coupled receptor, is necessary for glioblastoma growth. Oncogenesis 5, e263.

Bayin, N.S., Frenster, J.D., Sen, R., Si, S., Modrek, A.S., Galifianakis, N., Dolgalev, I., Ortenzi, V., Illa-Bochaca, I., Khahera, A., et al. (2017). Notch signaling regulates metabolic heterogeneity in glioblastoma stem cells. Oncotarget 8, 64932–64953.

Bohnekamp, J., and Schoneberg, T. (2011). Cell adhesion receptor GPR133 couples to Gs protein. J Biol Chem 286, 41912–41916.

Bruser, A., Schulz, A., Rothemund, S., Ricken, A., Calebiro, D., Kleinau, G., and Schoneberg, T. (2016). The Activation Mechanism of Glycoprotein Hormone Receptors with Implications in the Cause and Therapy of Endocrine Diseases. J Biol Chem 291, 508–520.

Cork, S.M., Kaur, B., Devi, N.S., Cooper, L., Saltz, J.H., Sandberg, E.M., Kaluz, S., and Van Meir, E.G. (2012). A proprotein convertase/MMP-14 proteolytic cascade releases a novel 40 kDa vasculostatin from tumor suppressor BAI1. Oncogene 31, 5144–5152.

Frenster, J.D., Inocencio, J.F., Xu, Z., Dhaliwal, J., Alghamdi, A., Zagzag, D., Bayin, N.S., and Placantonakis, D.G. (2017). GPR133 Promotes Glioblastoma Growth in Hypoxia. Neurosurgery 64, 177–181.

Frenster, J.D., Kader, M., Kamen, S., Sun, J., Chiriboga, L., Serrano, J., Bready, D., Golub, D., Ravn-Boess, N., Stephan, G., et al. (2020). Expression profiling of the adhesion G protein-coupled receptor GPR133 (ADGRD1) in glioma subtypes. Neurooncol Adv 2, vdaa053.

Frenster, J.D., and Placantonakis, D.G. (2018). Establishing Primary Human Glioblastoma Tumorsphere Cultures from Operative Specimens. Methods in molecular biology 1741, 63–69.

Gray, J.X., Haino, M., Roth, M.J., Maguire, J.E., Jensen, P.N., Yarme, A., Stetler-Stevenson, M.A., Siebenlist, U., and Kelly, K. (1996). CD97 is a processed, seven-transmembrane, heterodimeric receptor associated with inflammation. J Immunol 157, 5438–5447.

Hovelson, D.H., McDaniel, A.S., Cani, A.K., Johnson, B., Rhodes, K., Williams, P.D., Bandla, S., Bien, G., Choppa, P., Hyland, F., et al. (2015). Development and validation of a scalable next-generation sequencing system for assessing relevant somatic variants in solid tumors. Neoplasia 17, 385–399.

Hsiao, C.C., Chen, H.Y., Chang, G.W., and Lin, H.H. (2011). GPS autoproteolysis is required for CD97 to up-regulate the expression of N-cadherin that promotes homotypic cell-cell aggregation. FEBS Lett 585, 313–318.

Hsiao, C.C., Cheng, K.F., Chen, H.Y., Chou, Y.H., Stacey, M., Chang, G.W., and Lin, H.H. (2009). Site-specific N-glycosylation regulates the GPS auto-proteolysis of CD97. FEBS Lett 583, 3285–3290.

Hsiao, C.C., Keysselt, K., Chen, H.Y., Sittig, D., Hamann, J., Lin, H.H., and Aust, G. (2015). The Adhesion GPCR CD97/ADGRE5 inhibits apoptosis. Int J Biochem Cell Biol 65, 197–208.

Huang, Y.S., Chiang, N.Y., Hu, C.H., Hsiao, C.C., Cheng, K.F., Tsai, W.P., Yona, S., Stacey, M., Gordon, S., Chang, G.W., et al. (2012). Activation of myeloid cell-specific adhesion class G protein-coupled receptor EMR2 via ligation-induced translocation and interaction of receptor subunits in lipid raft microdomains. Mol Cell Biol 32, 1408–1420.

Iguchi, T., Sakata, K., Yoshizaki, K., Tago, K., Mizuno, N., and Itoh, H. (2008). Orphan G protein-coupled receptor GPR56 regulates neural progenitor cell migration via a G alpha 12/13 and Rho pathway. J Biol Chem 283, 14469–14478.

Kaur, B., Brat, D.J., Devi, N.S., and Van Meir, E.G. (2005). Vasculostatin, a proteolytic fragment of brain angiogenesis inhibitor 1, is an antiangiogenic and antitumorigenic factor. Oncogene 24, 3632–3642.

Kelley, L.A., Mezulis, S., Yates, C.M., Wass, M.N., and Sternberg, M.J. (2015). The Phyre2 web portal for protein modeling, prediction and analysis. Nature protocols 10, 845–858.

Kishore, A., Purcell, R.H., Nassiri-Toosi, Z., and Hall, R.A. (2016). Stalk-dependent and Stalk-independent Signaling by the Adhesion G Protein-coupled Receptors GPR56 (ADGRG1) and BAI1 (ADGRB1). J Biol Chem 291, 3385–3394.

Krasnoperov, V., Bittner, M.A., Holz, R.W., Chepurny, O., and Petrenko, A.G. (1999). Structural requirements for alpha-latrotoxin binding and alpha-latrotoxin-stimulated secretion. A study with calcium-independent receptor of alpha-latrotoxin (CIRL) deletion mutants. J Biol Chem 274, 3590–3596.

Krasnoperov, V., Lu, Y., Buryanovsky, L., Neubert, T.A., Ichtchenko, K., and Petrenko, A.G. (2002). Post-translational proteolytic processing of the calcium-independent receptor of alpha-latrotoxin (CIRL), a natural chimera of the cell adhesion protein and the G protein-coupled receptor. Role of the G protein-coupled receptor proteolysis site (GPS) motif. J Biol Chem 277, 46518–46526.

Langenhan, T. (2019). Adhesion G protein-coupled receptors-Candidate metabotropic mechanosensors and novel drug targets. Basic Clin Pharmacol Toxicol.

Lefkowitz, R.J., Cotecchia, S., Samama, P., and Costa, T. (1993). Constitutive activity of receptors coupled to guanine nucleotide regulatory proteins. Trends Pharmacol Sci 14, 303–307.

Liebscher, I., Schon, J., Petersen, S.C., Fischer, L., Auerbach, N., Demberg, L.M., Mogha, A., Coster, M., Simon, K.U., Rothemund, S., et al. (2014). A tethered agonist within the ectodomain activates the adhesion G protein-coupled receptors GPR126 and GPR133. Cell Rep 9, 2018–2026.

Liebscher, I., and Schoneberg, T. (2016). Tethered Agonism: A Common Activation Mechanism of Adhesion GPCRs. Handb Exp Pharmacol 234, 111–125.

Lin, H.H., Stacey, M., Yona, S., and Chang, G.W. (2010). GPS proteolytic cleavage of adhesion-GPCRs. Adv Exp Med Biol 706, 49–58.

Maser, R.L., and Calvet, J.P. (2020). Adhesion GPCRs as a paradigm for understanding polycystin-1G protein regulation. Cell Signal, 109637.

Mathiasen, S., Palmisano, T., Perry, N.A., Stoveken, H.M., Vizurraga, A., McEwen, D.P., Okashah, N., Langenhan, T., Inoue, A., Lambert, N.A., et al. (2020). G12/13 is activated by acute tethered agonist exposure in the adhesion GPCR ADGRL3. Nat Chem Biol.

Mehrotra, M., Duose, D.Y., Singh, R.R., Barkoh, B.A., Manekia, J., Harmon, M.A., Patel, K.P., Routbort, M.J., Medeiros, L.J., Wistuba, II, et al. (2017). Versatile ion S5XL sequencer for targeted next generation sequencing of solid tumors in a clinical laboratory. PLoS One 12, e0181968.

Morgan, R.K., Anderson, G.R., Arac, D., Aust, G., Balenga, N., Boucard, A., Bridges, J.P., Engel, F.B., Formstone, C.J., Glitsch, M.D., et al. (2019). The expanding functional roles and signaling mechanisms of adhesion G protein-coupled receptors. Ann N Y Acad Sci 1456, 5–25.

Moriguchi, T., Haraguchi, K., Ueda, N., Okada, M., Furuya, T., and Akiyama, T. (2004). DREG, a developmentally regulated G protein-coupled receptor containing two conserved proteolytic cleavage sites. Genes Cells 9, 549–560.

Okajima, D., Kudo, G., and Yokota, H. (2010). Brain-specific angiogenesis inhibitor 2 (BAI2) may be activated by proteolytic processing. J Recept Signal Transduct Res 30, 143–153.

Paavola, K.J., Stephenson, J.R., Ritter, S.L., Alter, S.P., and Hall, R.A. (2011). The N terminus of the adhesion G protein-coupled receptor GPR56 controls receptor signaling activity. The Journal of biological chemistry 286, 28914–28921.

Patra, C., van Amerongen, M.J., Ghosh, S., Ricciardi, F., Sajjad, A., Novoyatleva, T., Mogha, A., Monk, K.R., Muhlfeld, C., and Engel, F.B. (2013). Organ-specific function of adhesion G protein-coupled receptor GPR126 is domain-dependent. Proc Natl Acad Sci U S A 110, 16898–16903.

Petersen, S.C., Luo, R., Liebscher, I., Giera, S., Jeong, S.J., Mogha, A., Ghidinelli, M., Feltri, M.L., Schoneberg, T., Piao, X., et al. (2015). The adhesion GPCR GPR126 has distinct, domain-dependent functions in Schwann cell development mediated by interaction with laminin-211. Neuron 85, 755–769.

Promel, S., Frickenhaus, M., Hughes, S., Mestek, L., Staunton, D., Woollard, A., Vakonakis, I., Schoneberg, T., Schnabel, R., Russ, A.P., et al. (2012). The GPS motif is a molecular switch for bimodal activities of adhesion class G protein-coupled receptors. Cell Rep 2, 321–331.

Rath, A., Glibowicka, M., Nadeau, V.G., Chen, G., and Deber, C.M. (2009). Detergent binding explains anomalous SDS-PAGE migration of membrane proteins. Proc Natl Acad Sci U S A 106, 1760–1765.

Salzman, G.S., Zhang, S., Gupta, A., Koide, A., Koide, S., and Arac, D. (2017). Stachel-independent modulation of GPR56/ADGRG1 signaling by synthetic ligands directed to its extracellular region. Proc Natl Acad Sci U S A 114, 10095–10100.

Scholz, N., Monk, K.R., Kittel, R.J., and Langenhan, T. (2016). Adhesion GPCRs as a Putative Class of Metabotropic Mechanosensors. Handb Exp Pharmacol 234, 221–247.

Schoneberg, T., Kleinau, G., and Bruser, A. (2016). What are they waiting for?-Tethered agonism in G protein-coupled receptors. Pharmacol Res 108, 9–15.

Schulze, A., Kleinau, G., Neumann, S., Scheerer, P., Schoneberg, T., and Bruser, A. (2020). The intramolecular agonist is obligate for activation of glycoprotein hormone receptors. FASEB J 34, 11243–11256.

Stoveken, H.M., Hajduczok, A.G., Xu, L., and Tall, G.G. (2015). Adhesion G protein-coupled receptors are activated by exposure of a cryptic tethered agonist. Proceedings of the National Academy of Sciences of the United States of America 112, 6194–6199.

Vallon, M., and Essler, M. (2006). Proteolytically processed soluble tumor endothelial marker (TEM) 5 mediates endothelial cell survival during angiogenesis by linking integrin alpha(v)beta3 to glycosaminoglycans. J Biol Chem 281, 34179–34188.

Volynski, K.E., Silva, J.P., Lelianova, V.G., Atiqur Rahman, M., Hopkins, C., and Ushkaryov, Y.A. (2004). Latrophilin fragments behave as independent proteins that associate and signal on binding of LTX(N4C). EMBO J 23, 4423–4433.

Wilde, C., Fischer, L., Lede, V., Kirchberger, J., Rothemund, S., Schoneberg, T., and Liebscher, I. (2016). The constitutive activity of the adhesion GPCR GPR114/ADGRG5 is mediated by its tethered agonist. FASEB J 30, 666–673.

Yang, L.Y., Liu, X.F., Yang, Y., Yang, L.L., Liu, K.W., Tang, Y.B., Zhang, M., Tan, M.J., Cheng, S.M., Xu, Y.C., et al. (2017). Biochemical features of the adhesion G protein-coupled receptor CD97 related to its auto-proteolysis and HeLa cell attachment activities. Acta Pharmacol Sin 38, 56–68.

Zhu, B., Luo, R., Jin, P., Li, T., Oak, H.C., Giera, S., Monk, K.R., Lak, P., Shoichet, B.K., and Piao, X. (2019). GAIN domain-mediated cleavage is required for activation of G protein-coupled receptor 56 (GPR56) by its natural ligands and a small-molecule agonist. J Biol Chem 294, 19246–19254.

